# DNA replication fork stabilization and restart mechanisms revealed by biochemical reconstitution

**DOI:** 10.1101/2024.11.01.621508

**Authors:** Berta Canal, Agostina P. Bertolin, Giselle C. Lee, Masashi Minamino, John F.X. Diffley

**Affiliations:** Chromosome Replication Laboratory, The Francis Crick Institute, London, UK; Chromosome Segregation Laboratory, The Francis Crick Institute, London, UK

**Keywords:** DNA replication, DNA damage checkpoint, DNA polymerase α, Okazaki fragments, replication fork collapse, RPA depletion, PCNA, RFC, DNA polymerase δ, DNA polymerase ε

## Abstract

Understanding how DNA replication forks stall and restart and how the DNA damage checkpoint prevents irreversible fork collapse in molecular detail are crucial for understanding how cells maintain stable genomes and how they prevent the genetic instability that drives cancer. Here we describe the reconstitution of fork stalling and restart with purified budding yeast proteins. After nucleotide depletion, leading strand DNA synthesis quickly stops but CMG helicase continues to unwind and Okazaki fragments continue to initiate on the lagging strand. Incomplete Okazaki fragments sequester PCNA, RFC and DNA polymerases δ and ε which prevents normal DNA synthesis restart and exposes nascent DNA to nuclease attack. The DNA damage checkpoint limits this sequestration by restraining fork progression, which protects stalled forks from collapse and ensures restart.

## Introduction

Accurate and efficient DNA replication is crucial for genome maintenance. During chromosomal DNA replication, DNA unwinding by the CMG DNA helicase must be coupled with ongoing DNA synthesis^1–3^. Because of the anti-parallel nature of the DNA strands, DNA synthesis occurs continuously on the leading strand and discontinuously as Okazaki fragments (OkFs) on the lagging strand. OkFs are initiated by DNA polymerase α-primase (Polα): the DNA primase generates 7-10 nt RNA primers that are extended as DNA ∼20 nt by the DNA polymerase^4^. Rfc1-5 (RFC) loads the ring-shaped PCNA sliding clamp processivity factor on the 3’ DNA end generated by Polα, promoting completion of OkF synthesis by DNA polymerase δ (Polδ). Displacement of the 5’ end of the previous OkF by Polδ generates a flap which is cleaved by endonucleases, primarily Fen1, and the two OkFs are ligated together by ligase I (Lig1). Leading strands of the bidirectional replisomes are generated from OkFs; the leftward leading strand comes from the first OkF made by the rightward travelling replisome and vice versa^5–8^. When Polδ from these first OkFs reaches the CMG, DNA polymerase ε (Polε), which associates with CMG and is required for its formation, normally takes over for the duration of leading strand synthesis. Polε associates with dual processivity factors CMG helicase and PCNA to perform bulk leading strand synthesis^5–8^. We refer to the proteins required for OkF elongation (PCNA/RFC/Polδ), together with Polε, as the <u>Pr</u>ocessive DNA <u>S</u>ynthesis <u>M</u>achinery (PrSM), that together promote fast, processive and accurate DNA synthesis of leading and lagging strands.

DNA replication forks can stall as a result of nucleotide depletion or at DNA lesions from endogenous or exogenous sources. In response to fork stalling, cells activate the DNA damage checkpoint pathway that promotes a coordinated cellular response to ensure cell survival^9–14^. Fork stalling in the absence of the checkpoint results in replication fork collapse, accumulation of single-stranded (ssDNA), DNA damage, and cell death^15–22^. Restoration of a functional checkpoint after fork stalling does not restore cell viability, suggesting that the checkpoint prevents a catastrophic, irreversible event associated with stalled forks^16,18^. We currently do not fully understand what happens when replication forks stall, how replication forks restart or how the DNA damage checkpoint prevents irreversible fork collapse. In this paper we use DNA replication with purified proteins to characterise fork stalling, restart, and the mechanism of fork stabilization by the DNA damage checkpoint.

## Results

### DNA replication fork stalling and restart in vitro

We exploited the reconstitution of DNA replication with purified budding yeast proteins^5,23^ on a circular 10.6 kb DNA template containing a single origin of replication^5^ to study the nature of DNA replication fork stalling and restart. Replication of this template at our standard deoxynucleoside triphosphate (dNTP) concentration (80 μM each dNTP) in the absence of Fen1 or ligase results in production of leading strand products of approximately half the plasmid length (∼5.3 kb), and similar levels of DNA synthesis as discontinuous Okazaki fragments (OkFs) in the lagging strand (∼0.2-0.4 kb) within 10 min (**Figure S1A**). To induce DNA replication fork stalling, we performed DNA replication at several reduced dNTP concentrations (**Figure S1B**). At 1 µM total dNTP concentration (the ‘low dNTP concentration’ throughout the manuscript), leading strand DNA synthesis is initially slow (∼0.12 kb/min) and then stops after 6-8 min (**Figures 1A** and **S1C**). OkFs initiate but remain short (<0.1 kb); they primarily comprise DNA not RNA because they have incorporated α^32^P-dCTP and are resistant to alkaline hydrolysis during the processing of samples. OkFs continue to initiate and accumulate even after leading strand synthesis stops, resulting in 2-3X more incorporation in the lagging strand than in the leading strand after 30 min (**Figures 1A** and **1B**). Therefore, CMG continues to unwind DNA, leading strand synthesis stops, but OkFs continue to initiate and accumulate.

**Figure 1.**
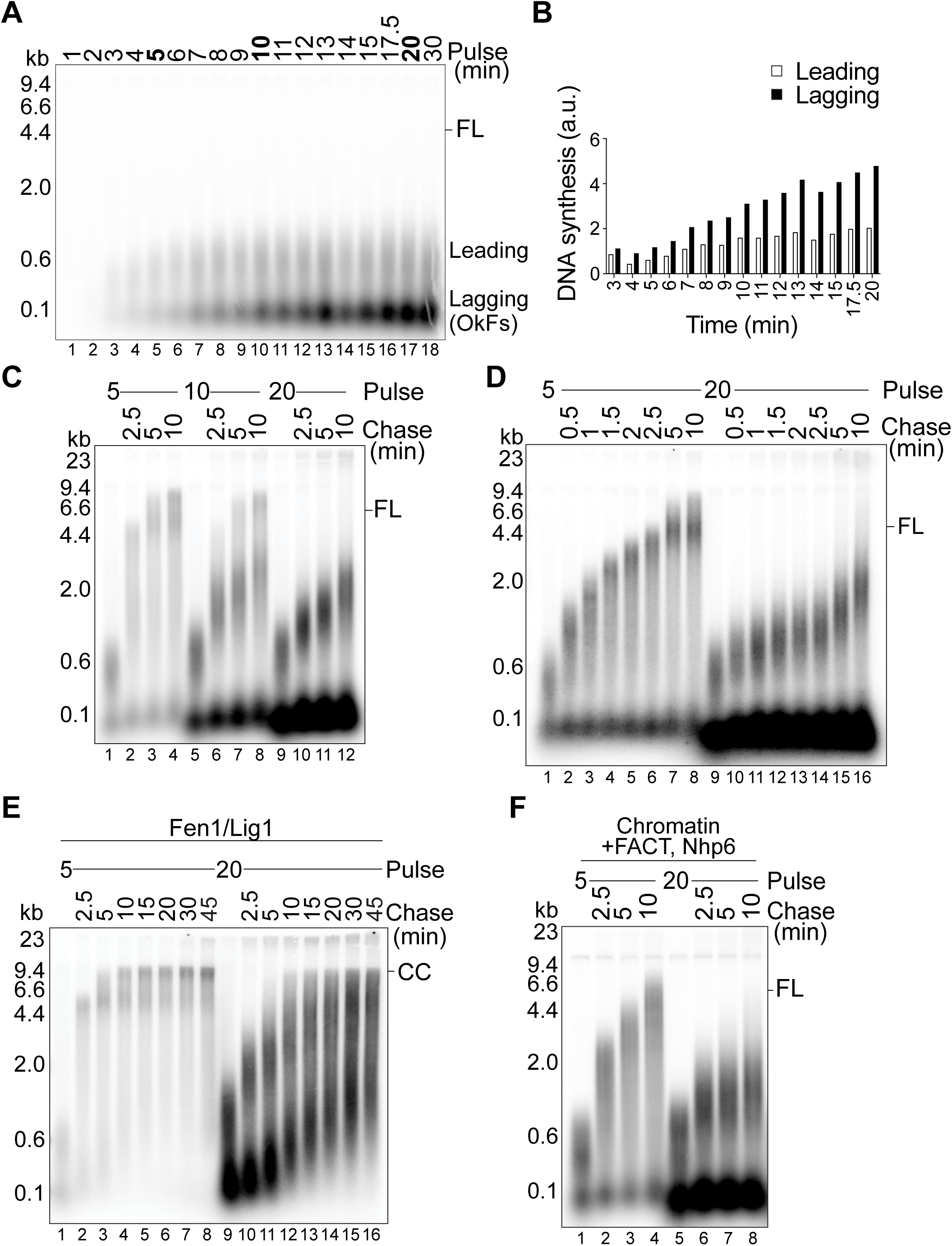
DNA replication fork stalling and restart in vitro. **(A)** Time course of DNA replication reaction at 1 μM (low) dNTPs with exo^−^ Polε. **(B)** Quantification of leading and lagging strand signal intensities from **A**. **(C)** Pulse-chase DNA replication reactions with WT Polε. 1 μM (low) dNTPs pulse for 5, 10 and 20 min chased each with 600 μM unlabelled dNTPs for 2.5, 5 and 10 min. <u>Unless specified, all subsequent</u> <u>pulse-chase reactions followed this same experimental strategy and were done with WT</u> <u>Polε.</u> **(D)** Pulse-chase DNA replication reactions. 5 and 20 min low dNTP pulses chased each at 0.5 min intervals, plus 5 and 10 min. **(E)** Pulse-chase DNA replication reaction always in the presence of Fen1 and Lig1. **(F)** Pulse-chase DNA replication reactions on a chromatin template. 5 and 20 min low dNTP pulses chased each for 2.5, 5 and 10 min. (FL) represents Full leading strand Length.

We added a high concentration of unlabelled dNTPs after different times at low dNTP concentration to promote and visualize DNA synthesis restart from stalled forks. After 5 min at low dNTP concentration, the amount of leading and lagging strand synthesis is roughly equivalent and synthesis is still occurring slowly (**Figure 1A**); dNTP addition at this point induced rapid synthesis, reaching near full length by 2.5 min (**Figure 1C**). However, after 20 min at low dNTP concentration, when leading strand synthesis has been stalled for >10 min and excess initiated OkFs have accumulated (**Figure 1A**), addition of dNTPs led to leading strand synthesis restart that was very slow and far from complete after 10 min (**Figure 1C**) and even after 60 min (**Figure S1D**). After 10 min at low dNTP concentration, two distinct populations of restart were observed, one fast and one slow (**Figure 1C**, lanes 6-8). This indicates that the difference in restart between 5 and 20 min is not a progressive slowing of restarting forks, but likely due to the acquisition of a discrete mode of slow DNA synthesis after extended fork stalling. Similar results were obtained when replication products were resolved on formaldehyde/formamide denaturing gels without alkaline treatment (**Figure S1E**), indicating that the apparent aberrant restart is not due to alkaline hydrolysis of the nascent DNA at positions of incorporated ribonucleotides, which could have been favoured after extended incubation at low dNTP concentration. Restart speed after 5 min at low dNTP concentration is comparable to unperturbed replication (**Figures 1D**, **S1F** and Yeeles *et al*^5^), and was unaffected by omission of Polδ (**Figure S1G**), indicating that Polε is sufficient for leading strand synthesis during short stalling and restart. Restart after 20 min was roughly 5 times slower than after 5 min (**Figure 1D** and **S1F**). Excess OkF initiation after 20 min and aberrant restart were still seen when we used a catalytically inactive Polε mutant (ΔCAT-Polε)^5^ and omitted Polδ from reactions (**Figure S1H**), indicating that DNA synthesis during the aberrant restart is catalysed primarily by Polα.

Inclusion of Fen1 and Lig1, enzymes that promote OkF maturation and ligation, during stalling and restart led to production of a completely replicated closed circular plasmid (CC) after restart of the 5 min incubation (**Figure 1E**). However, it did not restore normal leading strand synthesis restart after 20 min at low dNTP concentration and most replication products never reached completion (CC). Though there was some ligation of OkFs evident during restart, this was only partial even after 45 min. Therefore, the presence of OkF maturation machinery during stalling did not suppress aberrant restart. Note that during the low dNTP pulse, no ligation was observed with Fen1 and Lig1 at 5 min at low dNTP concentration indicating that little or no OkF completion and maturation occurred even from the beginning of the reaction.

Replication at low dNTP concentration of a chromatinised DNA template^24^ was very similar to naked DNA (**Figure 1F**): leading strand synthesis stopped at a simlar point and excess, short OkFs continued to accumulate over time. Therefore, the core DNA replication machinery can promote continued unwinding through chromatin even when leading strand DNA synthesis is inhibited. Similar to naked DNA, addition of dNTPs after 5 min at low dNTP concentration promoted efficient leading strand synthesis restart whereas restart was aberrant after 20 min, indicating that the presence of chromatin does not rescue defects in replication fork restart.

### The DNA damage checkpoint protects stalled forks

The checkpoint kinase Rad53 slows replication forks by phosphorylating and inactivating Mrc1 and Mcm10^25^. Omission of Mrc1 and reduction of Mcm10 concentration (Mcm10 cannot be omitted because it is required for initiation), slows replication fork progression^26–29^ and prevented accumulation of OkFs at later times, which allowed normal restart after 20 min at low dNTP concentration (**Figure 2A**). Addition of Rad53-phosphorylated Mrc1 and Mcm10 during fork stalling followed by unphosphorylated Mrc1 and Mcm10 during restart, which mimics events during fork stalling and restart *in vivo*, also limited OkF generation during fork stalling and rescued restart (**Figures 2B** and **S2**). Therefore, the Rad53 checkpoint kinase can protect stalled DNA replication forks at least in part by preventing unrestrained DNA unwinding.

**Figure 2.**
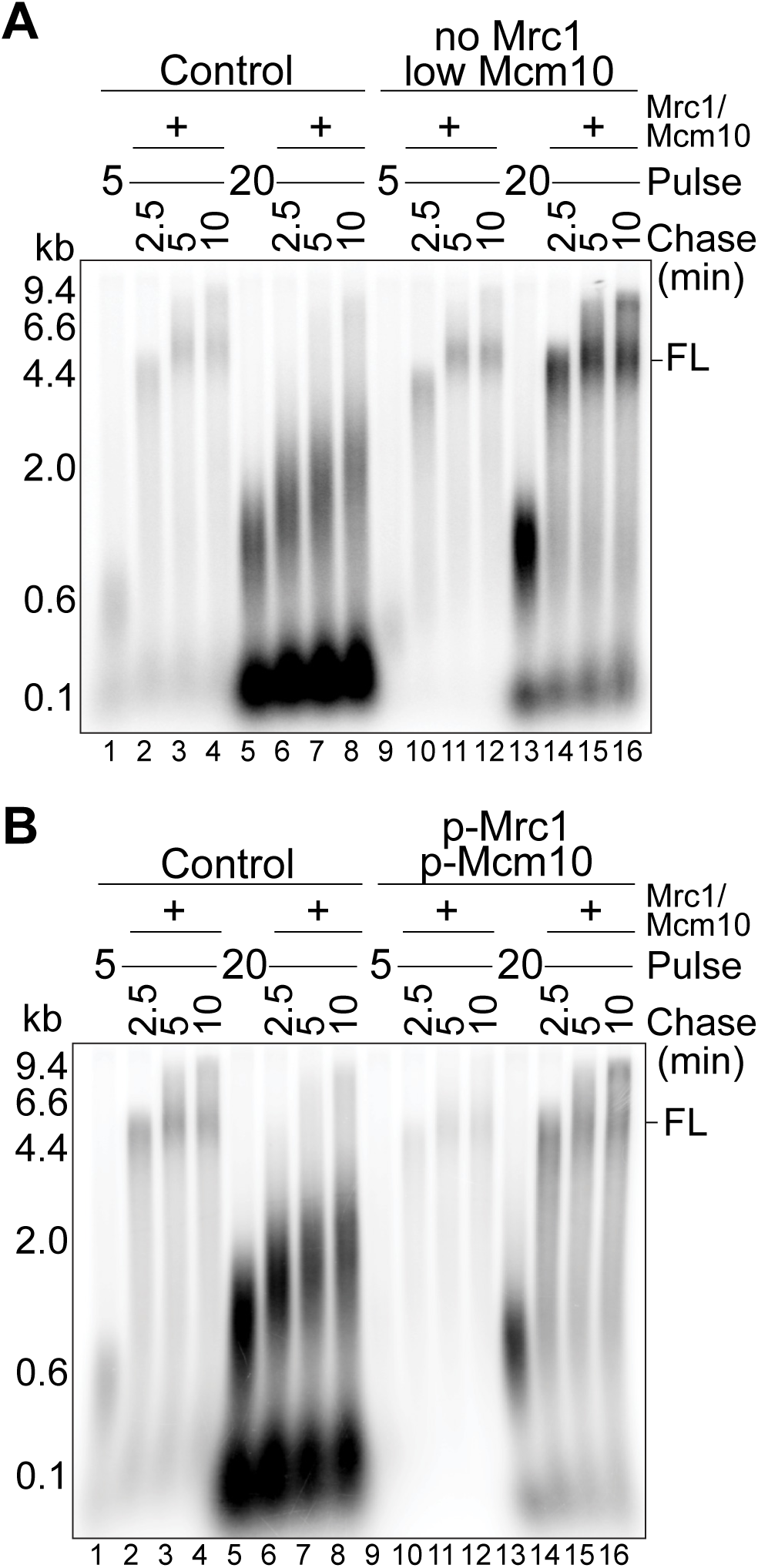
The DNA damage checkpoint protects stalled forks. **(A)** Pulse-chase DNA replication reactions. Reactions contained 20 nM Mrc1 and Mcm10 (Control) or no Mrc1 and 5 nM (low) Mcm10. 30 sec before chasing for 2.5, 5 and 10 min, 20 nM Mrc1 were added to all pulses. **(B)** As in **A**, but with Control or Rad53-phosphorylated 20 nM Mrc1 and Mcm10 (p-Mrc1/p-Mcm10). 30 sec before chasing for 2.5, 5 and 10 min, 60 nM unphosphorylated Mrc1 was added to all pulses. (FL) represents Full leading strand Length.

### Excess OkF initiation prevents restart

How does unrestrained fork progression prevent normal restart after 20 min of stalling? Generation of excess ssDNA could deplete soluble RPA and affect restart, as has been postulated as a cause of fork collapse in human cells lacking a functional checkpoint^30^. Whilst 80 nM RPA is sufficient for maximal levels of DNA replication in the absence of stalling (**Figure 3A**) even 10 times more RPA (800 nM RPA) did not rescue fork restart after either 20 or 10 min at low dNTP concentration (**Figures 3B** and **S3A**), arguing that RPA depletion is unlikely to be the cause of the aberrant restart observed upon fork stalling in our system.

**Figure 3.**
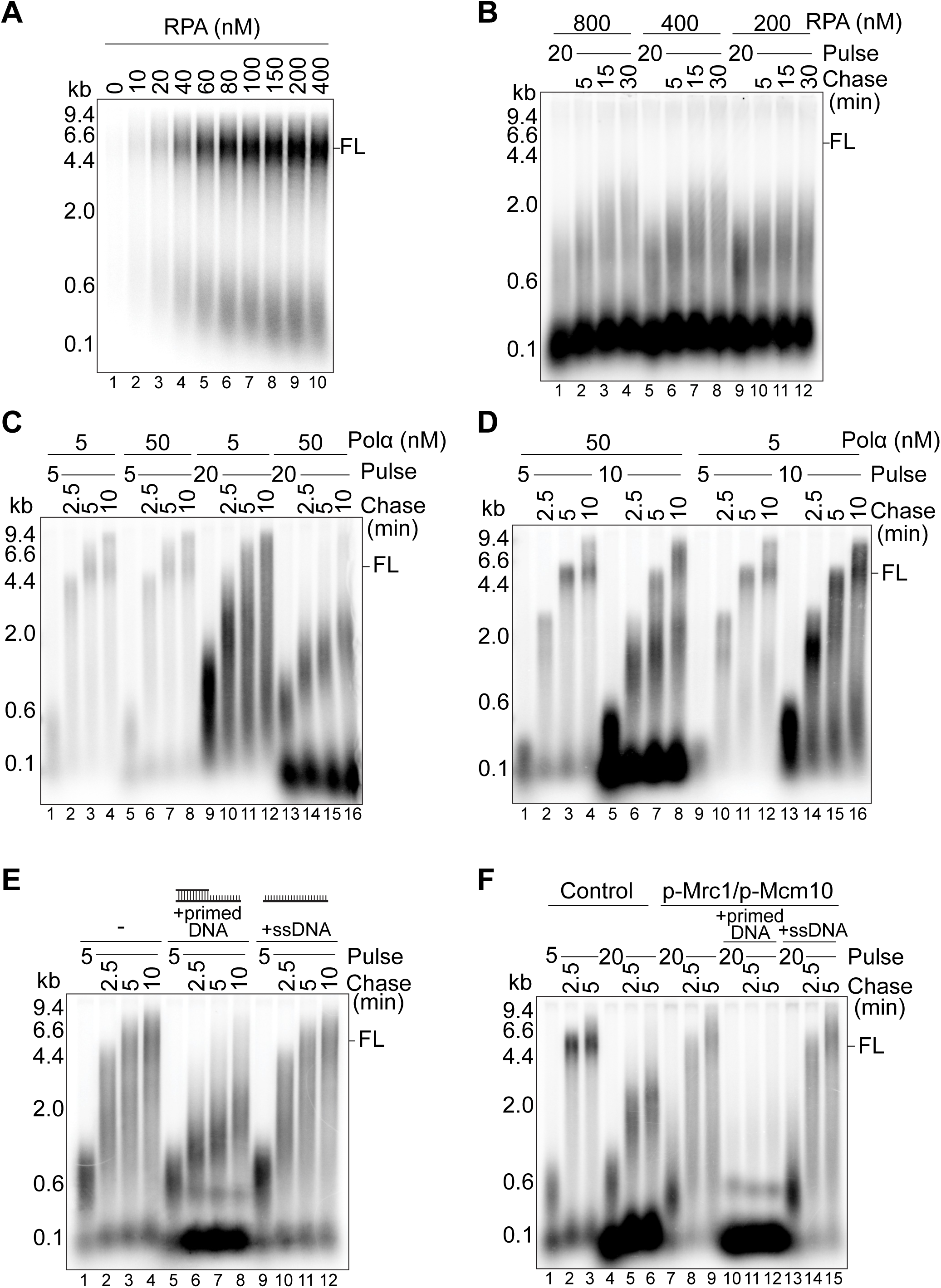
Excess OkF initiation prevents restart. **(A)** DNA replication for 7.5 min with 80 μM dNTPs at increasing RPA concentrations. **(B)** Pulse-chase DNA replication reactions. 20 min pulses with 800, 400 or 200 nM RPA, chased each for 5, 15 and 30 min. **(C)** Pulse-chase DNA replication. 5 and 20 min pulses with 5 or 50 nM Polα, chased each for 2.5, 5 and 10 min. **(D)** Pulse-chase DNA replication reactions. 5 and 10 min low dNTP pulses with 50 nM (standard) or 5 nM (low) Polα, chased each for 2.5, 5 and 10 min. **(E)** Pulse-chase DNA replication with 40 nM exo^−^ Polε. Buffer, 100 nM competitor primed DNA or ssDNA were added 4 min into the 5 min pulse, and chased each for 2.5, 5 and 10 min. **(F)** Pulse-chase DNA replication reactions containing 20 nM Mrc1 and Mcm10 (Control) or Rad53-phosphorylated 20 nM p-Mrc1/ 20 nM p-Mcm10. Buffer, 100 nM competitor primed DNA or ssDNA were added at 5 min into the 20 min pulse. 30 sec before chasing for 2.5, 5 and 10 min, 60 nM unphosphorylated Mrc1 was added to all pulses. (FL) represents Full leading strand Length.

We next asked whether continued OkF initiation and accumulation during extended CMG unwinding (**Figures 1A** and **1B**) was responsible for the aberrant restart. The frequency of OkF initiation on naked DNA templates can be modulated by changing Polα concentration^5^. Reducing Polα concentration during low dNTP treatment partially rescued the restart defect after 20 min (**Figure 3C**) and completely rescued restart after 10 min at low dNTP concentration (**Figure 3D**). Further reduction in Polα level resulted in reduced overall replication, presumably because of defects in establishing leading strand as well as lagging strand replication, and was therefore not interpretable.

As an alternative approach, we asked whether addition of competitor DNA mimicking either ssDNA or a short incomplete OkF could prevent normal restart after 5 min at low dNTP concentration (**Figure S3B**). Addition of a a 61 nt ssDNA oligonucleotide (competitor ssDNA) had no effect on restart after 5 min at low dNTP concentration, whilst the same oligonucleotide with an anealed 30 nt complement (competitor primed DNA) blocked normal restart after 5 min at low dNTP concentration (**Figure 3E**) resulting in restart very similar to that seen after 20 min. Addition of the same OkF-mimic also caused aberrant restart after 20 min with phosphorylated Mrc1 and Mcm10 (**Figure 3F**). Taken together, these results indicate that the continued accumulation of OkFs rather than ssDNA is the cause of the aberrant restart seen after 20 min at low dNTP concentration, and the checkpoint acts to prevent accumulation of excess OkFs.

### Depletion of PrSM factors contributes to fork stalling and causes aberrant restart

Excess incomplete OkFs could sequester any or all PrSM factors and, consistent with this, more PCNA was loaded onto DNA after 20 min than 5 min at low dNTP concentration (**Figures S4A**-**S4C**). Moreover, addition of extra PCNA, RFC, Polε and Polδ together, rescued restart from forks stalled for 20 min at low dNTP concentration (**Figure 4A**). Individual dropouts showed that all four PrSM factors contributed to efficient restart after 20 min (**Figures 4B** and **4C**), indicating that each PrSM factor is likely depleted to some extent. PCNA and RFC by themselves improved restart (**Figure 4B**) and dropout of either PCNA or RFC led to reduced restart (**Figure 4C**). Although dropout of Polε had a smaller effect than dropout of Polδ, there was a very substantial reduction in rescue when both polymerases were omitted indicating that, in the absence of extra Polδ, extra Polε can support restart (**Figure 4B**).

**Figure 4.**
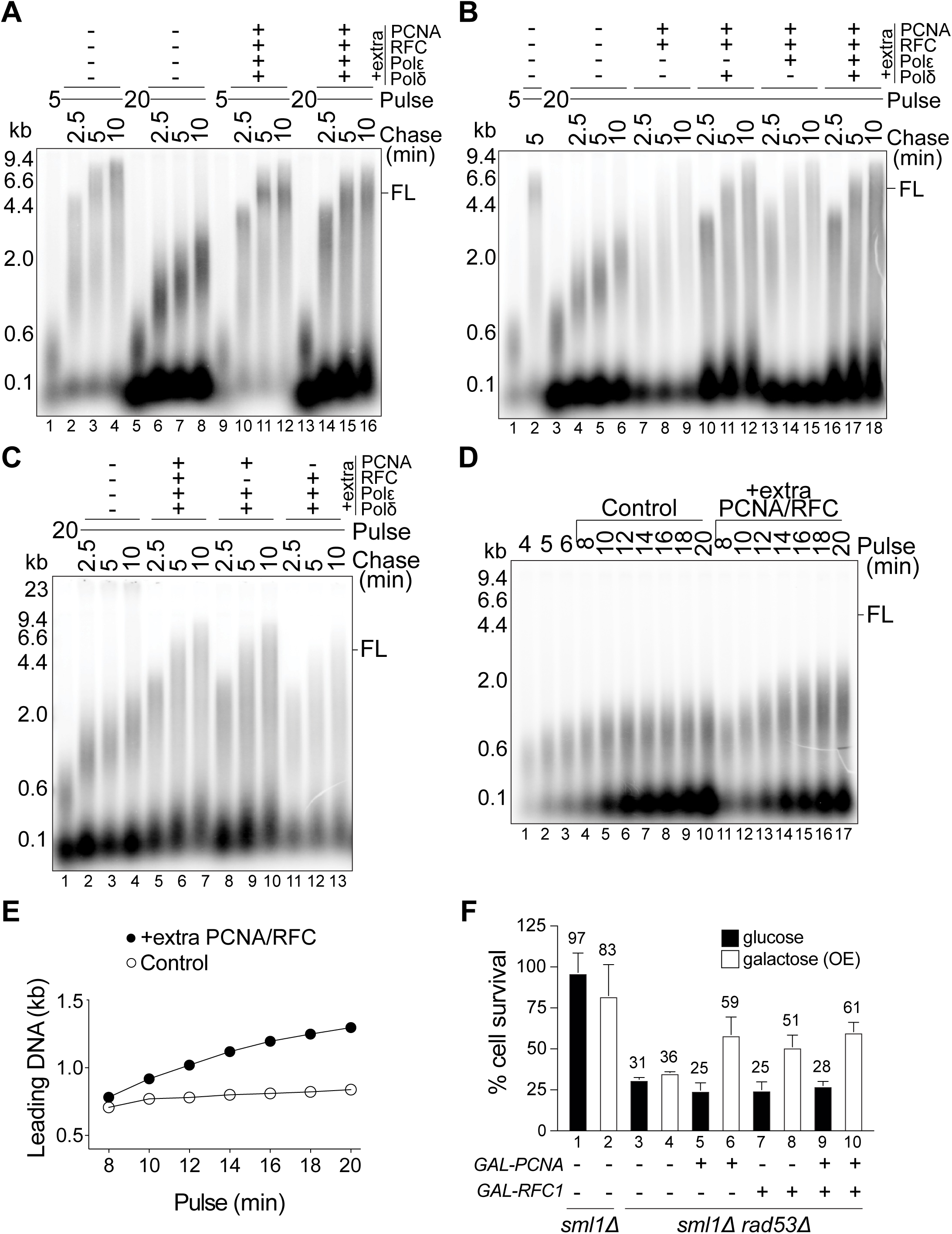
Depletion of PrSM factors contributes to fork stalling and causes aberrant restart. **(A)** Pulse-chase DNA replications. 5 and 20 min pulses. 30 sec before chasing, the buffers (−) or 250 nM PCNA, 125 nM RFC, 100 nM WT Polε and 10 nM Polδ were added to each pulse. **(B)** As in **A**, different PrSM factor combinations. **(C)** As in **A**, different PrSM factor combinations. **(D)** Time course of DNA replication reaction at low dNTP concentrations with WT Polε. Samples were taken at 4, 5 and 6 min before reaction was divided in two. At 7 min, either the buffers (Control) or the “extra PCNA/RFC” mix containing 250 nM PCNA and 125 nM RFC were added to each reaction. Time course samples were then taken from each reaction every 2 min for 20 min. **(E)** Quantification of bulk leading strand synthesis in “Control” and “extra PCNA/RFC” reactions from **D**. **(F)** Cell viability assay. Lanes 1-2: percentage of *sml1*Δ cells surviving after 1h in 0.01% MMS-containing YP-glucose (black) and YP-galactose (white), relative to survival in α-factor. Lanes 3-10: percentage survival in MMS of *sml1*Δ*rad53*Δ cells overexpressing none, PCNA, Rfc1 or PCNA+Rfc1, relative to their survival in α-factor and to survival of *sml1*Δ cells in MMS. Average % survival shown above bars. Error bars represent standard deviation. (FL) represents Full leading strand Length.

DNA synthesis stopped in the leading strand while lagging strand synthesis continued (**Figure 1A**), indicating that leading strand stalling occurred despite the continued presence of dNTPs. During the low dNTP incubation, we found that addition of extra PCNA and RFC promoted further leading strand synthesis without additional dNTPs (**Figures 4D** and **4E**). Therefore, the concentration of PCNA and RFC that is required and sufficient to sustain leading strand synthesis at the beginning of the reaction at low dNTP concentration, becomes insufficient as DNA replication proceeds at low dNTP concentration, indicating that replication at low dNTP concentration leads to early depletion of PrSM factors and contribute to fork stalling. Depletion of PCNA has relatively mild effects on leading strand synthesis during replication at high dNTP concentrations^5^; however, omission of PCNA from the beginning of the reaction at low dNTP concentration profoundly affected leading strand DNA synthesis (**Figure S4D** and Yeeles *et al*^5^). This residual synthesis is catalysed by Polα since similar synthesis is seen when PCNA and Polδ are both omitted in the presence of either WT Polε or ΔCAT Polε (**Figure S4E**). Taken together, these experiments indicate that PrSM depletion contributes to fork stalling at low dNTP concentration in addition to being critical for restart.

Mrc1 phosphomimicking mutants that reduce DNA replication fork speed have been shown to partially suppress the DNA damage sensitivity of a *rad53* mutant *in vivo*^25^. We wanted to determine whether excess PrSM factors could also suppress the effects of fork stalling in a checkpoint-deficient background. To avoid documented cell lethality associated with chronic overexpression of PCNA^31^, we designed an experimental strategy to induce an acute overexpression of PCNA and/or Rfc1 from a galactose-inducible promoter together with entry into S phase in the presence of methyl methanesulfonate (MMS). Under these conditions, the sensitivity of a *rad53* mutant to MMS was significantly decreased by exression of Rfc1 and PCNA individually or in combination (**Figures 4F**, **S4F** and **S4G**), consistent with the idea that PrSM depletion contributes to cell lethality in the absence of a functional DNA damage checkpoint.

### PrSM factors protect nascent DNA from nuclease attack

DNA synthesis at low dNTP concentration progressed with similar dynamics using WT or exonuclease mutant (exo^−^) Polε up to 20 min; and restart was similar with both polymerases (**Figure S5A**). However, the signal intensity and leading and lagging strand lengths were reduced at later times in reactions containing WT but not exo^−^ Polε (**Figures 5A**, **S5A** and **S5B**), indicating that, at later times at low dNTP concentration, the exonuclease of Polε was degrading nascent DNA.

**Figure 5.**
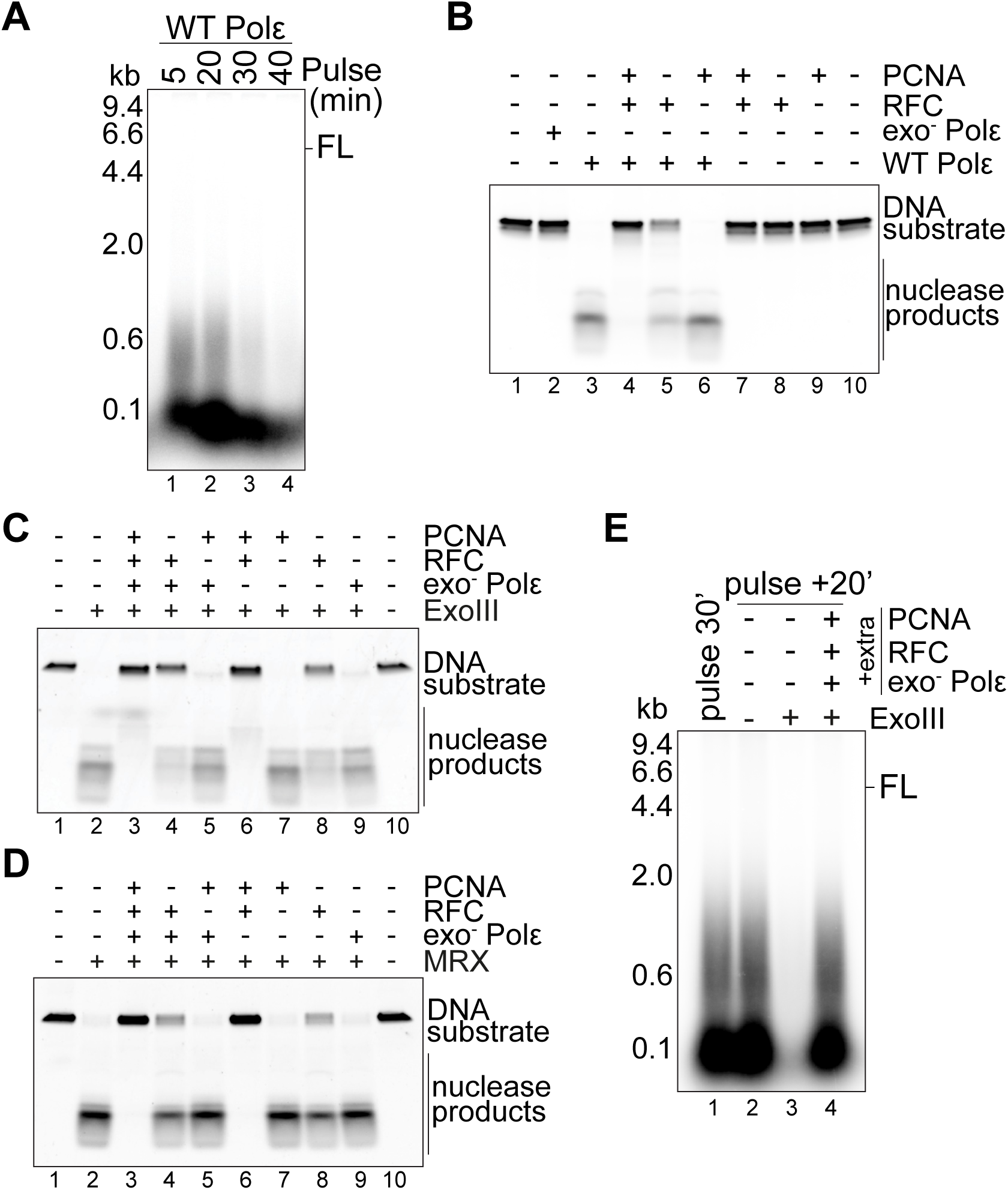
PrSM factors protect nascent DNA from nuclease attack. **(A)** Time course of DNA replication reaction in low dNTPs with WT Polε. **(B)** Nuclease reactions (5 min) with 10 nM exonuclease mutant (exo^−^) or WT Polε in the absence or presence of 125 nM RFC with or without 250 nM PCNA separated in TBE-PAGE and imaged for Cy3 fluorescence. **(C)** Nuclease reactions (5 min) with 2U of ExoIII nuclease (NEB) in the presence of different PrSM factor combinations containing 250 nM PCNA, 125 nM RFC, 100 nM exo^−^ Polε or their buffers. **(D)** As in **C**, but with 100 nM MRX nuclease for 45 min. **(E)** DNA replication reaction in low dNTPs with exo^−^ Polε for 30 min (pulse, lane 1). Pulse sample is then divided (lanes 2-4) and incubated for 20 min at 30°C with buffer or 20U ExoIII nuclease alone or in the presence of extra 250 nM PCNA, 125 nM RFC and 100 nM exo^−^ Polε. (FL) represents Full leading strand Length.

This suggested that depletion of PrSM factors may make DNA ends available for nucleolytic attack. To test this, we used a primed DNA model substrate (**Figure S5C**). The exonuclease of Polε degrades the substrate effectively, and RFC with or without PCNA provides protection from DNA degradation by Polε (**Figure 5B**). We next asked whether PrSM factors could protect DNA from other 3’ to 5’ exonucleases, the bacterial ExoIII and the yeast Mre11-Rad50-Xrs2 (MRX) complex (**Figures S5D** and **S5E**). **Figures 5C** and **5D** show that all PrSM factors contribute to maximum protection from both nucleases. Drop-outs of PCNA or RFC resulted in reduced protection (lanes 4 and 5). RFC provided protection on its own (lane 8). Exo^−^ Polε provided only very weak protection in the absence of PCNA and RFC (lane 9) but in their presence contributed to maximum protection (**Figure S5F**).

We then asked whether PrSM factors protect nascent DNA from degradation by nucleases during replication. Addition of the ExoIII nuclease after 30 min at low dNTP concentration resulted in degradation of leading and lagging nascent DNA products, which was prevented by addition of extra PrSM factors (**Figure 5E**). Therefore, PrSM depletion, in addition to affecting DNA synthesis restart, exposes nascent DNA ends, making them susceptible to nucleolytic attack. Of the four PrSM factors, RFC is especially good at protecting exposed DNA ends from nuclease attack.

## Discussion

From our work using a defined, reconstituted DNA replication system a picture emerges of 1) what happens when DNA replication forks stall in response to nucleotide depletion, 2) how forks restart and 3) how the DNA damage checkpoint protects stalled forks (**Figure 6**). Leading strand synthesis at 1 µM dNTP stalls after ∼5 min and ∼0.6 kb of synthesis but OkF synthesis continues for at least another 15 min resulting in ∼3X more incorporation. Based on the amount of synthesis (3X ∼0.6 = ∼1.8 kb) and the size of the OkFs (∼0.1 kb), after 20 min at low dNTP concentration, there are approximately 18 OkFs for every leading strand. From the start, these OkFs are incomplete, since even at the 5 min time point, inclusion of Fen1 and Lig1 did not lead to OkF ligation (**Figure 1E**). CMG unwinds DNA at a rate of ∼0.14 kb/min in the absence of DNA synthesis and the presence of Mrc1^25^; therefore, not more than 2.7 kb of DNA would be unwound in 20 min; consequently, the 18 OkFs are initiated roughly every 150 nt or less and are, therefore, more densely packed along the lagging strand than at 80 µM dNTPs, where the OkFs are ∼300 nt (**Figure S1A**). Consistent with this, OkFs only increase in size to ∼0.1 kb even after normal restart (**Figure 1C**). Therefore, OkF initiation after uncoupling from leading strand synthesis occurs frequently along the template at low dNTP concentration.

**Figure 6.**
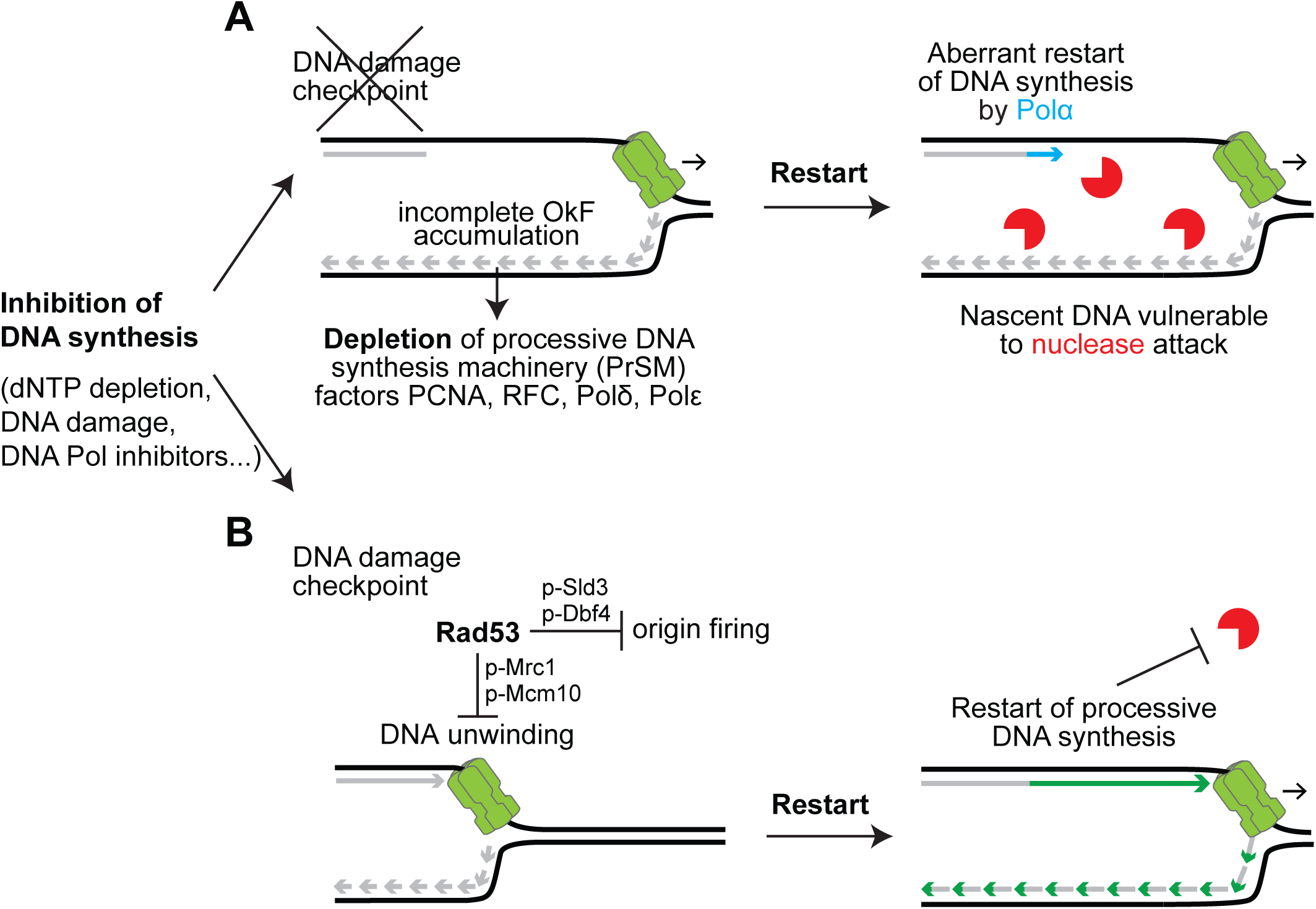
Mechanisms of checkpoint-dependent fork stabilization and restart. Inhibition of DNA synthesis causes fork stalling. When fork stalling occurs in the absence of the checkpoint (**A**), DNA unwinding continues and allows excess generation of incomplete OkFs, which sequester the PrSM factors. PrSM depletion causes aberrant restart by Polα and exposes nascent DNA to nuclease attack. When fork stalling occurs in the presence of the checkpoint (**B**), the Rad53 DNA damage checkpoint kinase prevents excess OkF generation by phosphorylating the replication proteins Mrc1, Mcm10, Sld3 and Dbf4 to inhibit DNA unwinding and origin firing. By preventing accumulation of incomplete OkFs, the checkpoint prevents PrSM depletion, which allows processive DNA synthesis restart and protects nascent DNA from nuclease attack.

Our results provide information about how DNA synthesis restarts after stalling. After a short (5 min) time at low dNTP concentration, leading strand synthesis resumes normally even in the absence of Polδ (**Figure S1G**) suggesting that Polε remains engaged with the 3’ end of the leading strand even as leading strand synthesis is slowing. We note that although Polε does not require PCNA for DNA synthesis in high dNTPs^5^, at low dNTP concentration PCNA loading by RFC is essential for leading strand DNA synthesis and restart by Polε (Yeeles *et al*^5^ and **Figures 4D**, **S4D** and **S4E**). The overall rate of leading strand synthesis at 80 µM dNTPs is greatly stimulated by PCNA^5^ suggesting transient stalling or uncoupling might occur without PCNA even at normal dNTP concentrations. Lagging strand synthesis by Polδ also appeared to restart normally after 5 min at low dNTP concentration as we saw near complete ligation of OkFs after adding dNTPs (**Figure 1E**).

After extended time (10-20 min) at low dNTP concentration, normal restart no longer occurs after dNTP addition. Instead, leading strand synthesis rate after restart is very slow. Polα is sufficient for this synthesis rate, and even after extended incubation, lagging strands are not completely ligated (**Figure 1E**) indicating that lagging strand synthesis restart is also aberrant. This restart resembles restart in human cells after aphidicolin arrest with checkpoint inhibitors described in the accompanying manuscript in that the rate of replication after restart is very slow and completely dependent upon Polα. Our results suggest that RPA depletion is not the cause of this phenomenon, consistent with work in the accompanying manuscript showing that aberrant fork restart precedes RPA exhaustion. Low levels of RPA (**Figure 3A**) reduce the frequency of OkF initiation without affecting the leading strand synthesis rate, consistent with previous results using a more minimal replication system^23^. At very low RPA levels, the amount but not the length of the leading strand is diminished (**Figure 3A**), suggesting there is a defect in generating the Okazaki fragment that seeds leading strand synthesis, but once initiated, leading strand synthesis is minimally affected by RPA depletion. In the accompanying manuscript, we show that RPA exhaustion contributes to DNA damage primarily at late times. Thus, RPA depletion may inhibit OkF synthesis at late times, contributing to more ssDNA and further RPA depletion. Aberrant restart is due to the sequestration of PrSM factors by excess OkFs, which prevents resumption of processive DNA synthesis (**Figure 6**). In contrast to leading strand synthesis restart after 5 min at low dNTP concentration, for which Polε is sufficient, Polδ appears to be more important than Polε in restart after 20 min (**Figures S1F** and **4B**). We suggest that this is because Polδ synthesises DNA rapidly and processively with PCNA outside the context of CMG and thus is the ideal polymerase to ‘recouple’ leading strand synthesis to CMG^5^, in much the same way it establishes the leading strand from an OkF during the early stages of replication^5,7,8^ and how it recouples leading strand synthesis to CMG after translesion synthesis^32^.

Our model is supported by observations *in vivo* using strand-specific ChIP-Seq methods showing that the DNA damage checkpoint prevents PCNA accumulation on hydroxyurea-stalled lagging strands^33^ and uncoupling of leading and lagging strand synthesis^34^; moreover, these phenotypes are suppressed by Mrc1 deletion^35^. Although this was attributed to accumulation of toxic ssDNA^36^, we propose that this is instead due to accumulation of excess OkFs. Our results are also consistent with the fact that Polδ mutants show synthetic lethality with the checkpoint kinases Rad53 and Mec1 in yeast^37^ and even transient depletion of Polδ causes lethality of Mec1 mutant yeast cells^38^. Similarly, tumour cells with compromised Polδ activity were shown to have higher vulnerability to cancer therapies using checkpoint inhibitors^39^.

We propose that, by preventing PrSM depletion, the checkpoint protects nascent DNA ends from enzymatic attack by exonucleases and potentially other enzymes such as helicases. PrSM factors protect 3’ DNA ends from nuclease attack, both from heterologous nucleases like ExoIII and potentially relevant nucleases like Mre11 and the proofreading exonuclease of Polε (**Figure 5**), the latter of which has been implicated in fork collapse in yeast in the absence of a functional checkpoint^40^. The requirements for protection and restart are not identical; RFC by itself is sufficient for some protection but not for restart, and thus its depletion may have the most profound consequences *in vivo*, as argued in the accompanying manuscript. A recent report^41^ indicates that RFC can bind to 5’ as well as 3’ DNA ends which could also protect nascent DNA from attack by 5’ to 3’ nucleases like Exo1^42,43^. Regardless, PrSM factors may also indirectly protect DNA ends by promoting OkF completion and ligation. The DNA damage checkpoint modulates the enzymatic activity of several nucleases directly^40,42,44,45^ which also contributes to protection of nascent DNA ends.

We do not know how much of the irreversible loss of viability upon fork stalling in the absence of the checkpoint is due to enzymatic attack of nascent DNA facilitated by PrSM depletion and how much is due to the irreversible loss of processive leading and lagging strand synthesis after PrSM depletion in yeast. Loss of the Exo1 nuclease prevented degradation of stalled forks and improved cell viability in certain backgrounds^18,43,46^ and the exonuclease activity of Polε has been implicated in cell death in checkpoint mutants^40^. In our reconstituted system, loss of processive DNA synthesis after PrSM depletion was irreversible unless new PrSM factors were added to the reaction. Irreversibility was still observed when we added Elg1-Rfc2-5, the PCNA unloader (**Figures S6A**-**S6B**) even after pre-phosphorylation by Rad53^33^ (**Figure S6C**). This is not very surprising for three reasons: first, excess OkF initiation depletes all four PrSM factors, not just PCNA. Second, after unloading PCNA, there still remains an excess of 3’ ends from incomplete OkFs competing with the 3’ end of the leading strand. And third, Elg1-Rfc2-5 was proposed to unload PCNA after OkF maturation^47^, but maturation does not occur at low dNTP concentration after addition of Lig1 and Fen1 (**Figure 1E**), presumably because they cannot act on incomplete OkFs and they require interaction with PCNA, and so may also be compromised after PrSM depletion. Further work is needed to understand how PCNA loading and unloading is regulated in unperturbed and stalled forks.

Given these uncertainties, it appears that prevention of PrSM depletion is the best strategy for survival, and our results indicate that the DNA damage checkpoint plays a crucial role in preventing PrSM depletion by slowing unwinding from existing forks through phosphorylation and inhibition of Mrc1 and Mcm10. Prevention of continued origin firing through Rad53-phosphorylation of Sld3 and Dbf4 likely also contributes to preventing excessive OkF synthesis and PrSM depletion^48,49^.

## Materials and Methods

### Plasmids and yeast strains

Yeast strains generated in this study are listed in **Table S1**. Full sequence maps of the plasmids generated in this study can be found in **Table S2**.

To perform cell viability assays, strains were derived from diploid W303 strain containing SML1 and RAD53 gene deletions and transformed with plasmids pBC65 (Gal-PCNA) and pBC66 (Gal-3xFlag-Rfc1).

To express recombinant MRX complex, yBC107 strain was constructed on the background yeast strain yJF1^50^ by sequentially integrating plasmid pBC42 (Rad50) linearised with NheI into HIS3 locus, and then plasmid pBC43 (Mre11 + Xrs2-TEV-3xFlag) linearised with StuI into URA3 locus.

To express recombinant Elg1-Rfc2-5 complex, yBC155 strain was constructed on the background yeast strain yAE36, previously used in the construction of Rfc1-5 expressing strain yAE41^5^, by integrating plasmid pBC52 (Elg1-3xFlag) linearised with NheI into HIS3 locus.

Strains for all other recombinant protein expression are as described^5,23^ except for Dpb11 that was expressed in yeast strain yVP8^25^, and Sld2 and RPA that were expressed in *E.coli* from plasmids pGC441^51^ and pJM126 (Addgene) respectively.

### Protein expression and purification

A summary of protein purification strategies used in this study can be found in **Table S3**. Proteins were essentially expressed and purified as described^52^ with the modifications detailed here. **S-CDK** purification included a MonoQ column prior to gel filtration. Protein was loaded and washed with 10 column volumes (CV) of 150 mM potassium acetate buffer and eluted with 20CV linear gradient 150-1000 mM potassium acetate before loading to Superdex 200. **Csm3/Tof1** was purified excluding the MonoQ column. **exo^−^ Polε** was purified like WT Polε except elution from heparin column was done with a linear gradient of 400-1500 mM potassium acetate. **ΔCAT Polε** was purified as described^5^. **Rad53** was purified as described^25^, using storage buffer containing 25 mM HEPES-KOH pH 7.6, 300 mM NaCl, 10% glycerol, 0.02% NP-40 and 1 mM DTT. **Sld2** was expressed in BL21-Codon Plus (DE3) - RIL competent cells from plasmid pGC441^51^. **FACT** and **Nhp6** were purified as described^24^. **Elg1-Rfc2-5** was expressed in budding yeast and purified like the Rfc1-5 complex (see **Table S1**, **S2** and **S3** for yeast strain, plasmid and expression details).

Untagged **RPA** was purified as described^53^ with the following modifications. BL21-Codon Plus (DE3) - RIL competent cells were transformed with plasmid pJM126 and grown overnight at 37°C without shaking in LB medium supplemented with 100 μg/ml ampicillin and 50 μg/ml chloramphenicol. In the morning, cells around OD 0.1-0.2 were shaked until OD 0.5 before adding 0.4 mM IPTG for 2h at 37°C. From here, all steps were performed in ice or at 4°C. Cells were lysed in RPA buffer (25 mM HEPES pH 7.5, 10% glycerol, 1 mM EDTA, 0.02% NP-40, 1 mM DTT) with 0.5 M NaCl (0.5M NaCl Buffer) supplemented with cOmplete™, EDTA-free Protease Inhibitor Cocktail (Sigma) doing 2 freeze-thaw cycles and sonication in ice-water for 5 min in cycles of 5 sec ON, 10 sec OFF at 35%. Lysate was first clarified at 45 krpm for 45 min at 4°C and then passed through a 0.22 uM filter. The clarified supernatant was applied to a 5 ml HiTrap® Blue High-Performance column connected to an AKTA Pure, and protein eluted sequentially with 1CV 0.5M NaCl Buffer, 1.5CV of 0.8M NaCl Buffer, 1.5CV of 0.5M NaSCN Buffer and 2CV 1.5M NaSCN Buffer. Pooled peak fractions were then loaded onto a 5 ml Bio-Scale Mini CHT Type I Cartridge (Biorad) in 10 mM NaH_2_PO_4_ Buffer and eluted sequentially with 2CV 40 mM NaH_2_PO_4_ Buffer, 120 mM NaH_2_PO_4_ Buffer and 500 mM NaH_2_PO_4_ Buffer. Most protein eluted in the 40 mM NaH_2_PO_4_ Buffer. Pooled peak fractions were dialyzed in 150 mM NaCl Buffer and loaded onto a 1 ml Mono Q column, washed with 20CV 150 mM NaCl Buffer, and eluted in a 20CV linear gradient from 150-1000 mM NaCl (RPA elutes ∼250 mM NaCl). Peak fractions were pooled and concentrated before loading onto a Superdex200 in 150 mM NaCl Buffer, after which peak fractions were pooled, concentrated, aliquoted and stored at −80°C.

**MRX** was expressed in budding yeast (see **Table S1** and **S2** for yeast expression strain and plasmid details). 8L cells were grown at 30°C to OD 1 in YP + 2% raffinose and protein expression was induced by addition of galactose to 2% for 2h. Cells were lysed with a Freezer/Mill in Buffer MRX (25 mM HEPES-KOH pH 7.6, 10% Glycerol, 1 mM EDTA, 0.02% NP-40-S and 1 mM DTT) with 500 mM NaCl (Buffer MRX 500) and protease inhibitors. All subsequent steps were conducted at 4°C. Lysate was cleared at 45,000 rpm for 45 min at 4°C. Cleared lysate and 2 ml Flag bead slurry were incubated 2h at 4°C before the resin was collected, and subsequently washed with 50 ml Buffer MRX 300, incubated/washed 10 min in Buffer MRX 300 with 10 mM Magnesium acetate and 0.55 mg/ml ATP and washed again with 50 ml Buffer MRX 300. MRX was eluted in 2 subsequent steps containing 0.5 mg/ml and 0.25 mg/ml Flag peptide in 5 ml Buffer MRX 300. The eluate was then slowly diluted 3-fold by addition of Buffer MRX (no salt) before loading to a 1 ml Heparin in Buffer MRX 100, washed with 10CV and eluted with linear gradient 100-1000 mM NaCl. Pooled peak fractions were concentrated to 0.4 ml and separated through a Superose6 gel filtration column in Buffer MRX 200. MRX containing fractions were pooled and concentrated to ∼ 0.5 mg/ml, aliquoted and stored at −80°C.

### DNA replication reaction with purified proteins

Replication reactions were performed as described in the original publication^5,52^. For every replication reaction, MCMs were loaded for 20 min at 30°C and 1250 rpm using 40 nM ORC, 40 nM Cdc6, 60 nM MCM-Cdt1, 4 nM 10.6kb ARS1-containing template DNA and 5 mM ATP in replication buffer adjusted to get final concentrations of 25 mM HEPES-KOH pH 7.6, 100 mM potassium glutamate, 10 mM magnesium acetate, 0.02% NP-40-S and 2 mM DTT (5 μl loading reactions). After 20 min of MCM loading, 50 nM DDK (in 5 μl) was added and incubated for 15 min. Then, in ice, a protein mix was added to give final concentrations of 40 nM Dpb11, 20 nM Polε, 20 nM GINS, 80 nM Cdc45, 20 nM CDK, 25 nM Sld3/7, 50 nM Sld2, 400 nM RPA, 20 nM Ctf4, 10 nM TopoI, 10 nM TopoII, 1 nM Polδ (unless specified), 25 nM PCNA, 30 nM Rfc1-5 (RFC), 20 nM Mrc1, 20 nM Csm3/Tof1, 50 nM Polɑ and 20 nM Mcm10 (10 μl replication reaction). In **Figure 1E**, 40 nM Fen1 and 40 nM Cdc9 (Lig1) were added. Finally, reactions were started by bringing the volume up to the total 10 μl with a ‘nucleotide’ mix containing 200 μM of each NTP (CTP, GTP, UTP), 1 μM (low) or 80 μM (control=high) dNTPs, and 10 nM (low) or 33 nM (control=high) ɑ^32^P-dCTP in replication buffer adjusted to increase final salt concentration to 225 mM potassium glutamate. Unless specified, all DNA replication reactions were done with WT Polε. pulse-chase reactions were initiated with 5, 10 or 20 min pulses containing 1 μM (low) dNTP and 10 nM ɑ^32^P-dCTP, and then each pulse was chased with 600 μM non-radioactive dNTPs for the specified times. ‘dNTP’ concentrations reflect the concentration of each dNTP (dCTP, dGTP, dTTP and dATP) in a mix. Reactions were stopped by the addition of 85 mM EDTA in ice, cleared from free nucleotides over an Illustra MicroSpin G-50 column, and denatured in sample buffer at final concentration of 2% sucrose, 0.02% BPB, 60 mM NaOH, 10 mM EDTA, separated on 0.8% alkaline agarose gels/running buffer containing 30 mM NaOH and 2 mM EDTA at approximately 1 volt/cm for 14-17h at room temperature, fixed in cold 5% trichloroacetic acid 30 min, dried on Whatman paper, exposed to phosphor screens, and scanned using a Typhoon phosphor imager.

For formaldehyde/formamide denaturing of DNA replication samples in **Figure S1E**, samples cleared in G-50 column were incubated in 0.3M NaCl for 2h at 55°C before denaturing DNA with final 9% formaldehyde and 50% formamide at 95°C for 5 min. Sample buffer was added containing final 20% glycerol, 1 mM EDTA pH 8.0 and 0.02% BPB/Xylene cyanole and then run in 0.8% agarose gels containing 6.66% formaldehyde and 1x MOPS-NaOH buffer pH 7 (20 mM MOPS free acid, 5 mM Na-Acetate anhydrous pH 5, 1 mM EDTA pH 8) with 1x MOPS-NaOH buffer pH 7 running buffer. Running conditions and gel processing were done as above.

For DNA replication on chromatin in **Figure 1F**, 50 μg of purified histone octamers was mixed with 50 μg of the 10.6 kb DNA template to a final volume of 200 μL in chromatin buffer (25 mM HEPES-KOH pH 7.6, 1 mM EDTA) containing 1 M NaCl. The histone-DNA mix was loaded into a D-TubeTM Dialyser Mini (Merck Millipore) and dialysed at 4°C against 0.5 L chromatin buffer of decreasing salt concentrations (1 M NaCl for 3 h, 0.75 M NaCl overnight, 0.5 M NaCl for 5 h, 0.0025 M NaCl overnight). After the final dialysis step, the dialysate is applied to a Superose 6 Increase 3.2/300 column (Cytiva) equilibrated in 25 mM HEPES-KOH pH 7.6, 150 mM NaCl. Peak fractions containing reconstituted chromatin were pooled and assessed by micrococcal nuclease digestion. Chromatin was dialysed against storage buffer (25 mM HEPES-KOH pH 7.6, 5 mM NaCl, 0.1 mM EDTA) and stored at 4°C.

For preparation of competitor DNAs used in **Figures 3E** and **3F**, the oligonucleotides (see **Figure S3B** and **Table S4** for details) were annealed to a final concentration of 5 μM each in 20 mM Tris–HCl pH 7.5, 5 mM magnesium acetate and 100 mM NaCl (100 μl reaction) by heating at 95°C for 2 min and letting cool down to 25°C. Competitor primed DNA was aliquoted and kept at −80°C. 0.5 μM competitor primed DNA or competitor ssDNA oligo was then incubated at with 2.5 μM Streptavidin (Invitrogen) in replication buffer (20 μl reactions) for 10 min in ice before adding to DNA replication reactions at final concentrations of 100 nM.

### Rad53 kinase reaction

For kinase reactions in **Figures 2B** and **3F**, Mrc1 and Mcm10 were incubated with Rad53 protein or buffer at 1:1:2 (Mrc1:Mcm10:Rad53) molar ratio in the presence of 5 mM ATP in replication buffer for 30 min prior to adding to 10 μl DNA replication reactions at final concentrations of 20 nM Mrc1 and 20 nM Mcm10.

For kinase reactions in **Figure S6C**, Elg1-Rfc2-5 and Rad53 or their buffers were incubated at 1:1 (Elg1-Rfc2-5:Rad53) molar ratio in the presence of 5 mM ATP in replication buffer for 30 min prior to adding to 10 μl DNA replication reactions at final concentration of 30 nM Elg1-Rfc2-5.

### Pull down of nascent DNA

DNA replication reaction (x10 reactions/condition, total 100 μl each) was performed as above but containing 100 nM RFC, 400 nM PCNA, and 1 μM dATP/dCTP/dGTP and (1:1) 7.5 μM dTTP and 7.5 μM Biotin-dUTP for 5 and 20 min. 600 μM dTTP were added at 5 min into the 20 min reaction to prevent/chase further incorporation of Biotin-dUTP to nascent DNA. Reactions were stopped by the addition of 85 mM EDTA in ice and cleared from free nucleotides over an Illustra MicroSpin G-50 column. At this point, 10% of volume was collected as “total” sample (5% for western blot and 5% for alkaline agarose electrophoresis). MagStrep “type 3” Strep-Tactin beads were pre-equilibrated in pull down buffer (25 mM HEPES-KOH pH 7.6, 50 mM NaCl, 2 mM EDTA, 0.02% NP-40-S and 2 mM DTT) and incubated with remaining sample for 30 min at 25°C and 1250 rpm. Beads were then bound to magnet, 10% of unbound volume was collected as “flow through” sample (and divided as above), and beads were washed 3 times with 100 μl pull down buffer. DNA was then eluted from beads incubating 15 min at 25°C and 1250 rpm in 20 μl elution buffer (25 mM Tris-Cl pH 7.5, 150 mM NaCl, 1 mM EDTA and 50 mM Biotin). Pull down sample was divided 80% for western blot and 20% for alkaline agarose electrophoresis. Samples for alkaline agarose gel electrophoresis were processed as above. Samples for western blot were denatured in SDS-DTT sample buffer, boiled, and separated in 10% TGX gel (BioRad) before blotting to nitrocellulose membrane. Membrane was blotted sequentially with anti-PCNA ([5E6/2], mouse monoclonal) and anti-Mcm6 (custom-made, rabbit polyclonal).

### Nuclease reaction

For DNA substrate preparation, oligonucleotides were annealed as for competitor DNA (see above) but to a final concentration of 0.5 μM each. See **Table S4** for oligonucleotide details.

Nuclease reactions (10 μl) were performed at 30°C and 1250 rpm in 100 mM K-glutamate DNA replication buffer. 10 nM annealed DNA substrate and 2.5 mM ATP were incubated with PCNA and RFC (or their buffers) for 5 min at 30°C before addition of the corresponding nuclease and, in **Figures 5C** and **5D**, also addition of the buffer or exo^−^ Polε. Nuclease reactions were incubated at 30°C for 5 min (Polε), 5 min (ExoIII) or 45 min (MRX). Reactions were then stopped by the addition of 5 μl 3x STOP buffer to have final concentrations of 30 mM EDTA, 0.5% SDS, 6.5% Ficoll and 0.3 mg/ml Proteinase K, and incubated at 30°C for 10 min before separating (dark) in a 20% native TBE-PAGE (Invitrogen) in 1xTBE running buffer at 150V for 75 min at room temperature, and Cy3 fluorescence was visualised with Amersham Imager 800, GE lifesciences.

### Cell viability assay

Cells were grown overnight at 30°C in YP + 2% raffinose, diluted in the morning in YP + 2% raffinose to OD 0.1 and grown to OD 0.3 before addition of 20 μg/ml ɑ-factor for 1h. At this point, cells were maintained in ɑ-factor for 1-2 more hours, but culture was divided in two and 2% galactose or 2% galactose was added to each half. A sample of G1-synchronised cells was taken, and cells were washed and released into YP + 2% galactose or 2% galactose + 0.01% Methyl Methanesulfonate (MMS) and 5 μg/ml nocodazole for 1h at 30°C before taking sample of MMS condition. To evaluate cell viability, 100 μl of cells were diluted (1:5000-1:20,000) in YP + 2% glucose and plated in YP + 2% glucose agar plates. Colonies on plates were left to grow for 2-3 days at 30°C and number of colonies/plate/condition were counted. For each strain and condition, we also collected samples for western blot and flow cytometry (see below).

### Western blot

For protein expression analysis, 1 ml cell culture from viability experiment in **Figure 4F** was extracted in 85% TCA and lysed for 40 sec at 6 m/s using a Fastprep. Extracts were then processed by 4-15% Tris-Glycerol SDS-PAGE, transferred to nitrocellulose, and immunoblotted with anti-PCNA ([5E6/2], mouse monoclonal), anti-Flag (Rfc1) ([M2], mouse monoclonal), anti-Rad53 ([Abcam ab104232], rabbit polyclonal) and anti-PGK ([Abcam 22C5D8], mouse monoclonal).

### Flow cytometry

For flow cytometric analysis, 200 μl cell culture form viability experiment in **Figure 4F** was fixed in 70% ethanol for 30 min and then treated overnight with 1 mg/ml RNAse A at 30 °C in 50 mM sodium citrate. Cells were stained with 4 μg/ml propidium iodide in 50 mM sodium citrate and analysed using a FACSCalibur (BD) and FlowJo Software.

### Quantification and statistical analysis

#### Quantification of DNA replication

DNA replication quantifications were performed using ImageJ software following signal image linearisation with ‘linearise gel data’ plug-in. Distance in the gel was calibrated to base pair length using the λ DNA-HindIII Digest ladder (NEB) with an exponential fit.

Quantification of DNA synthesis in leading and lagging strands in **Figure 1B** was done by plotting lane signal intensity and calculating the area under the curve for each strand using ImageJ.

Quantification of leading DNA replication length in **Figures 4E**, **S1C** and **S1F** were done using Graph-pad Prism by first fitting leading DNA synthesis intensity to a non-linear gaussian curve and calculating the mean (representing the maximum bulk DNA synthesis for each lane) and then, in **Figures S1C** and **S1F**, fitting the mean of each lane to a simple linear regression to calculate DNA synthesis speed.

#### Statistical analysis

Graphpad Prism software was used for all statistical analyses. Statistical methods are described in the figure legends as appropriate.

## Supplementary Tables

**Table S1.** Yeast strains generated in this study.

| Table S1. Yeast strains generated in this study. |  |
| --- | --- |
| Name | Genotype |
| yBC107<br>(OE coMRX) | *yJF1 + pBC42 (NheI)::HIS3 + pBC43 (StuI)::URA3* |
| yBC155<br>(OE coElg1-Rfc2-5) | *MATa ade2-1 ura3-1 his3-11,15 trp1-1 leu2-3,112 can1-100*<br>*bar1::Hyg*<br>*pep4::KanMX*<br>*ura3::URA3 pRS306-RFC2+CBP-RFC3*<br>*trp1::TRP1 pRS304-RFC4 + RFC5*<br>*his3::HIS3 pRS303-ELG1-3xFlag (pBC52 (NheI)::HIS3)* |
| yBC319<br>(sml1) | *W303-1a sml1Δ::KanMX* |
| yBC320<br>(rad53) | *W303-1a sml1Δ::KanMX rad53Δ::LEU2* |
| yBC296<br>(rad53 OE PCNA) | *W303-1a sml1Δ::KanMX rad53Δ::LEU2 + pBC65 (StuI)::URA3* |
| yBC297<br>(rad53 OE Rfc1) | *W303-1a sml1Δ::KanMX rad53Δ::LEU2 + pBC66 (BsgI)::TRP1* |
| yBC298<br>(rad53 OE<br>PCNA/Rfc1) | *W303-1a sml1Δ::KanMX rad53Δ::LEU2 + pBC65 (StuI)::URA3 +*<br>*pBC66 (BsgI)::TRP1* |

**Table S2.**
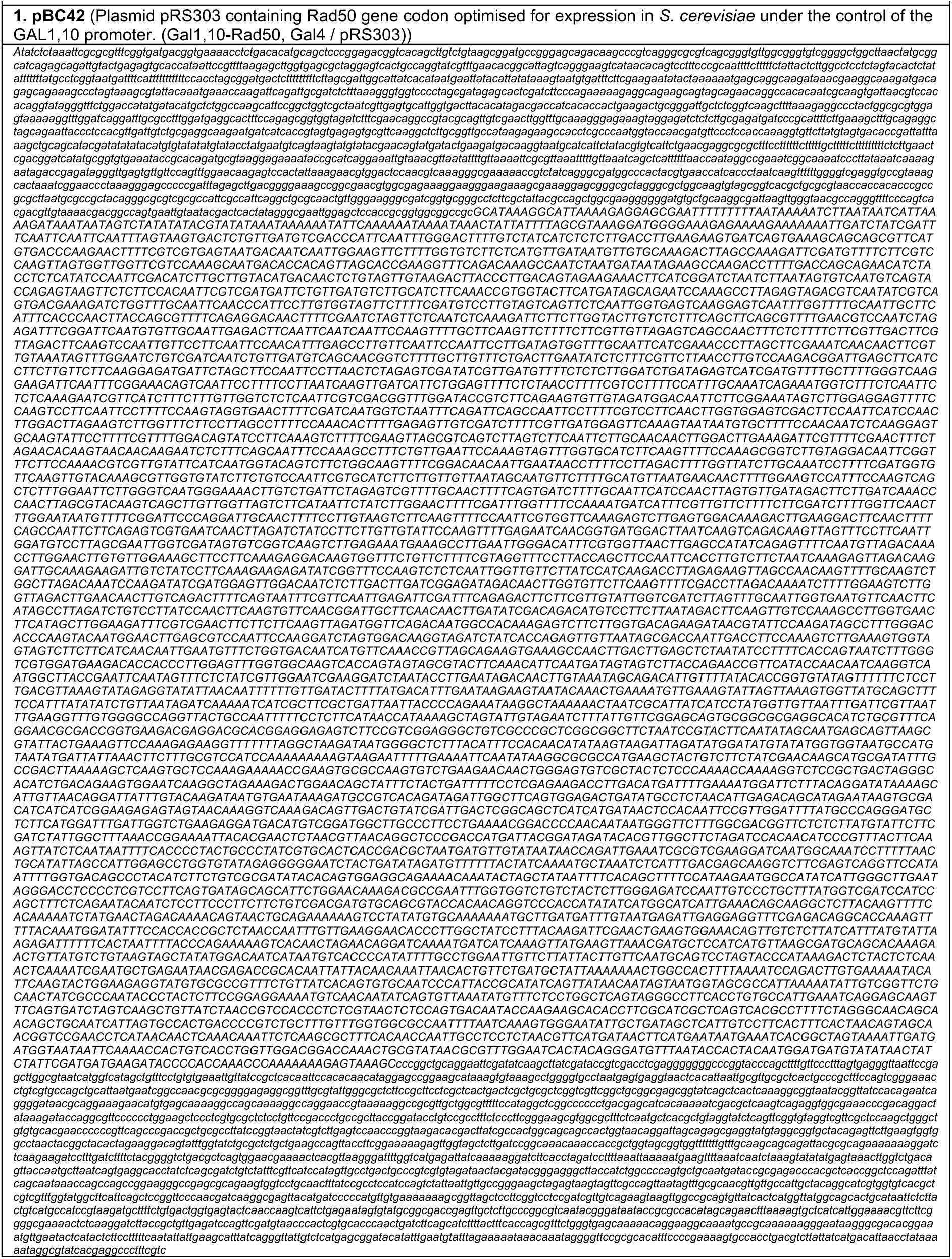
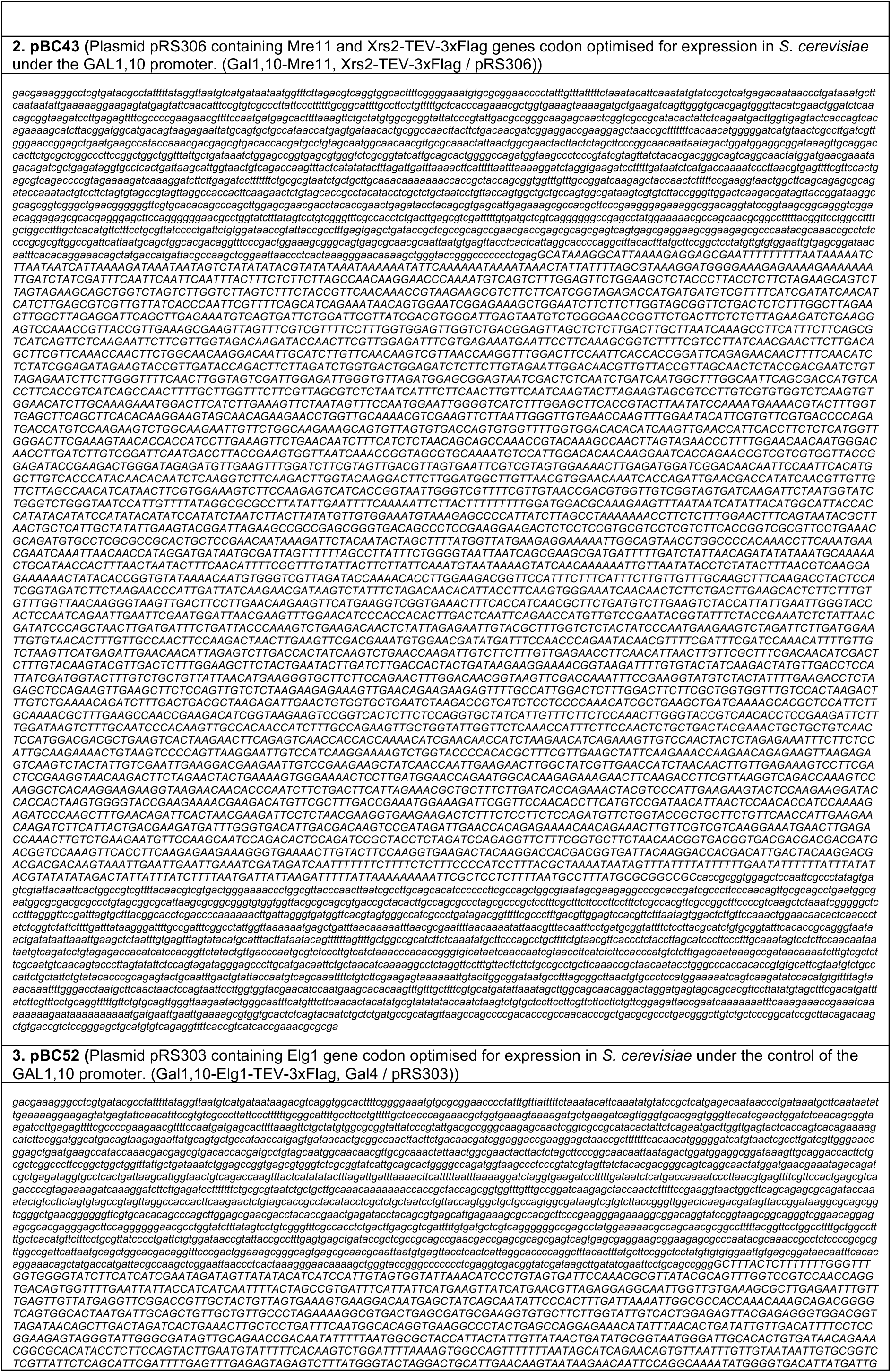
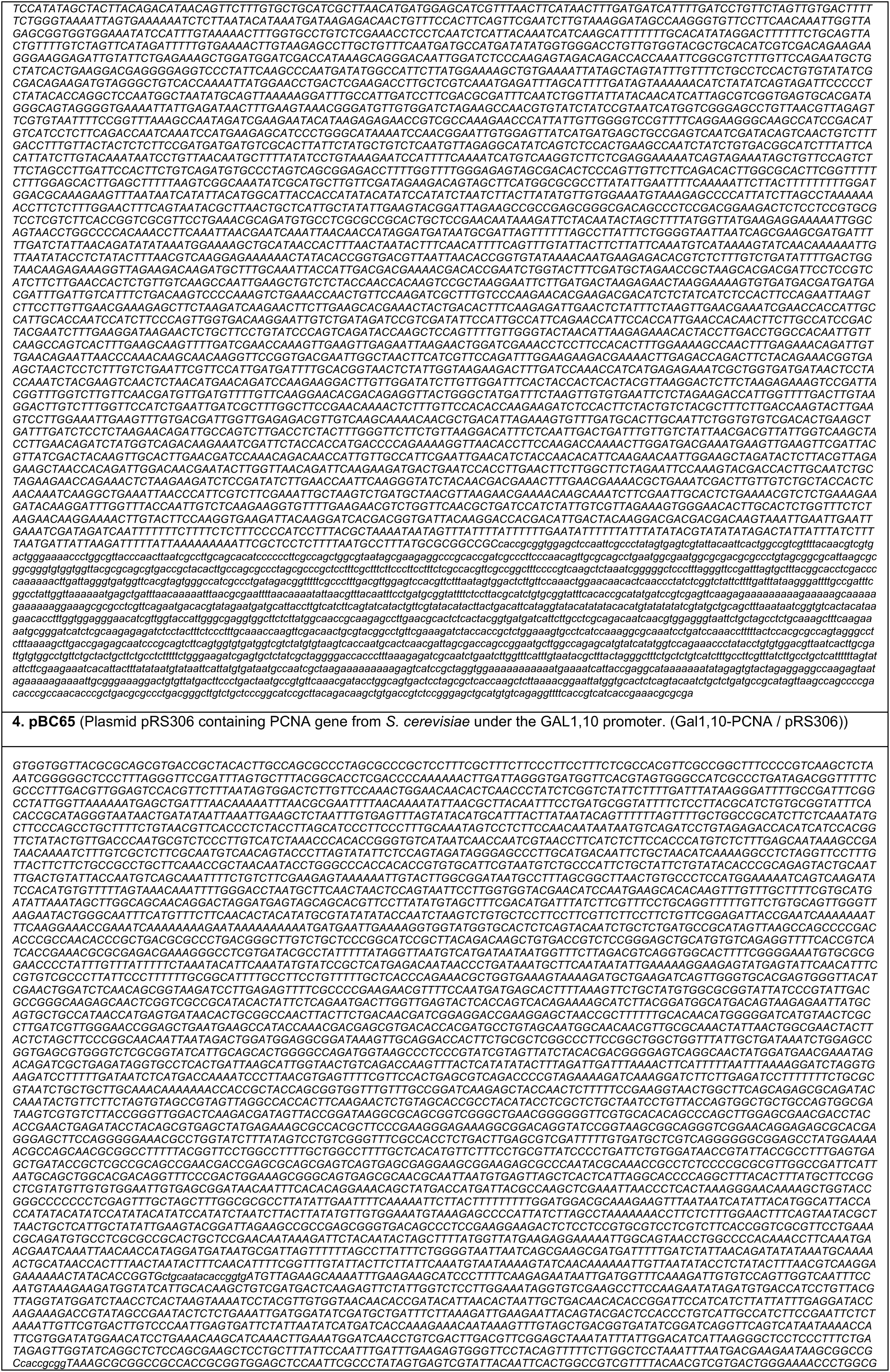
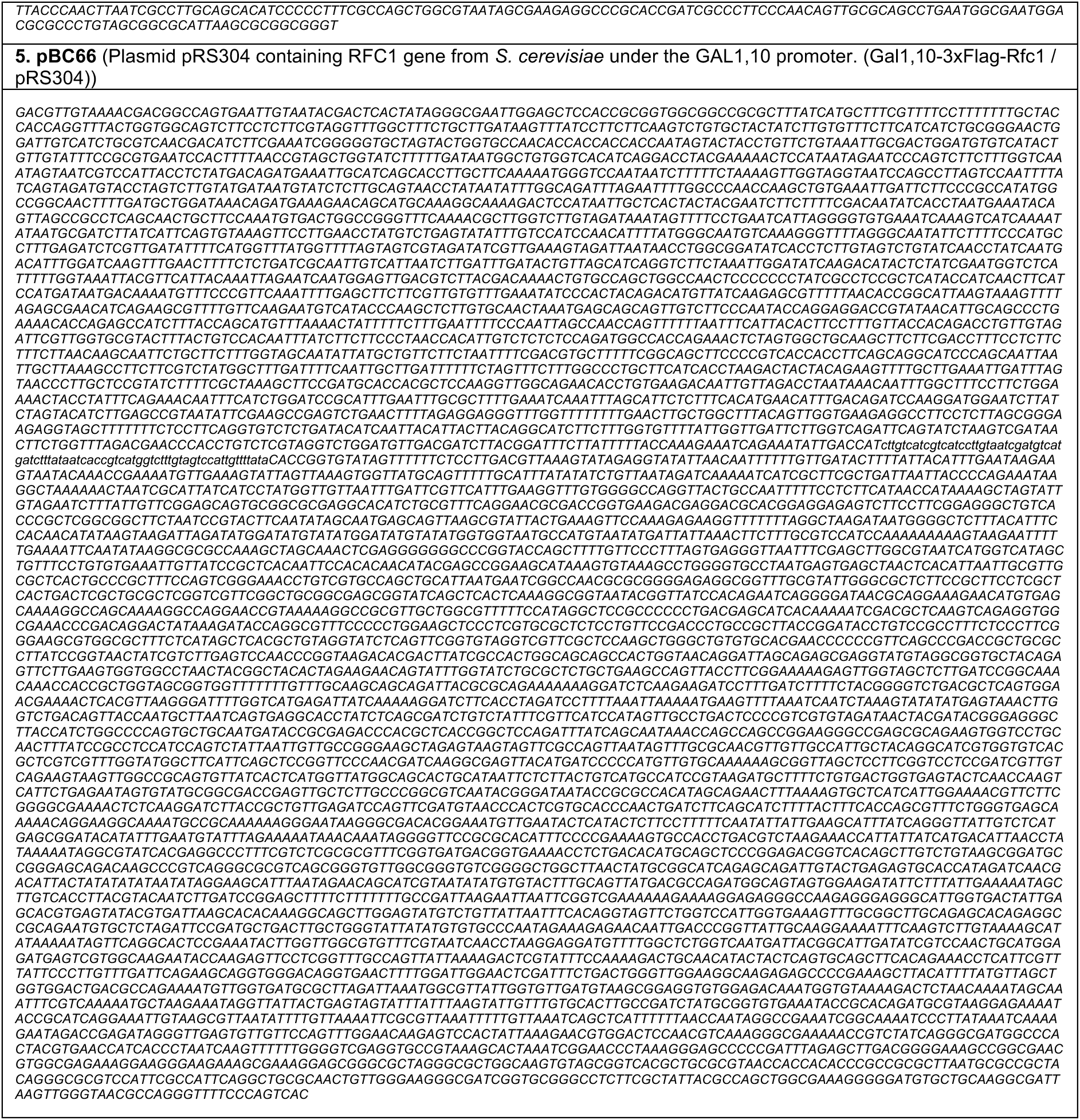
Plasmids generated in this study.

**Table S3.** Protein expression and purification strategies.

| **Protein** | **Expressio**<br>**n**<br>**organism** | **Pull down**<br>**strategy** | **N-/C- tag** | **Elution** | **Chromatography**<br>**columns** |
| --- | --- | --- | --- | --- | --- |
| **ORC** | *S. cerevisiae* | calmodulin | CBP-Orc1 | EGTA | HiLoad Superdex 16/600 |
| **Cdc6** | *E. coli* | glutathione | GST-Cdc6 | PreScission | Hydroxyapatite, dialysis |
| **MCM-Cdt1** | *S. cerevisiae* | calmodulin | CBP-Mcm3 | EGTA | HiLoad Superdex 16/600 |
| **DDK** | *S. cerevisiae* | calmodulin | CBP-Dbf4 | EGTA | Superdex200 |
| **Dpb11** | *S. cerevisiae* | 3xFlag | Dpb11-3xFlag | Flag peptide | Superdex200 |
| **Pol $\epsilon$** | *S. cerevisiae* | calmodulin | Dpb4-CBP | EGTA | Heparin, Superdex200 |
| **exo- Polε** | *S. cerevisiae* | calmodulin | Dpb4-CBP | EGTA | Heparin, Superdex200 |
| **ΔCAT Polε** | *S. cerevisiae* | calmodulin | Dpb4-CBP | EGTA | MonoQ, Superdex200 |
| **GINS** | *E. coli* | Ni-NTA | 6His-GINS | Imidazole | MonoQ, Superdex200 |
| **Cdc45** | *S. cerevisiae* | 3xFlag | internal 2xFlag | Flag peptide | Superdex200 |
| **CDK** | *S. cerevisiae* | calmodulin | CBP-Clb5 | EGTA | MonoQ, Superdex200 |
| **Sld3/Sld7** | *S. cerevisiae* | IgG<br>sepharose | TCP-Sld3 | TEV | NiNTA (remove TEV),<br>Superdex200 |
| **Sld2** | *E. coli* | glutathione | GST-Sld2 | PreScission | SP FF, dialysis |
| **RPA** | *E. coli* | - | - | - | HiTrap® Blue High<br>Performance, Bio-Scale<br>Mini CHT Type I<br>Cartridge, MonoQ,<br>Superdex200 |
| **Ctf4** | *S. cerevisiae* | calmodulin | CBP-Ctf4 | EGTA | MonoQ, Superdex200 |
| **TopI** | *S. cerevisiae* | calmodulin | CBP-TopI | EGTA | Superdex200 |
| **TopII** | *S. cerevisiae* | calmodulin | TopII-CBP | EGTA | MonoQ, Superdex200 |
| **Polδ** | *S. cerevisiae* | calmodulin | Pol32-CBP | EGTA | Heparin, Superdex200 |
| **Rfc1-RFC** | *S. cerevisiae* | calmodulin | CBP-Rfc3 | EGTA | SP FF, Superdex200 |
| **PCNA** | *E. coli* | ammonium<br>sulfate | - | - | Heparin/SP FF/DEAE,<br>MonoQ, Superdex200 |
| **Mrc1** | *S. cerevisiae* | 3xFlag | Mrc1-3xFlag | Flag peptide | MonoQ, Dialysis |
| **Csm3/Tof1** | *S. cerevisiae* | calmodulin | CBP-Csm3 | TEV | Superdex200 |
| **Polα** | *S. cerevisiae* | calmodulin | CBP-Pri1 | EGTA | MonoQ, Superdex200 |
| **Mcm10** | *E. coli* | Flag/NiNTA | His-Mcm10-Flag | Flag peptide/<br>Imidazole | NiNTA, Superdex200 |
| **Fen1** | *S. cerevisiae* | 3xFlag | Fen1-3xFlag | Flag peptide | Heparin, Dialysis |
| **Cdc9** | *S. cerevisiae* | 2xFlag | Cdc9-2xFlag | Flag peptide | MonoQ, Dialysis |
| **FACT** | *S. cerevisiae* | 3xFlag | 3xFlag-Pob3 | Flag peptide | - |
| **Nhp6** | *S. cerevisiae* | untagged | - | - | Negative TCA<br>precipitation, MonoS |
| **Rad53** | *E. coli* | NiNTA | Rad53-6His | Imidazole | Superdex200 |
| **MRX** | *S. cerevisiae* | 3xFlag | Xrs2-3xFlag | Flag peptide | Heparin, Superose6 |
| **Elg1-RFC** | *S. cerevisiae* | calmodulin | CBP-Rfc3 | EGTA | SP FF, Superdex200 |

**Table S4.**
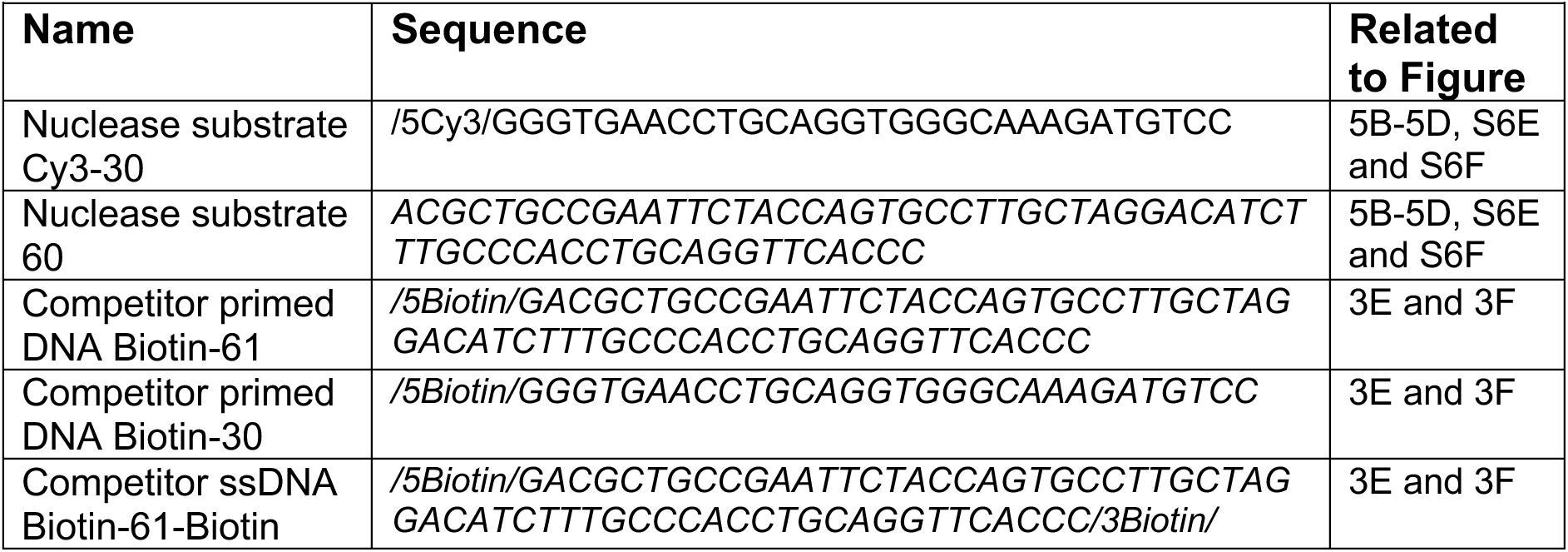
Oligonucleotides for DNA substrates.

## Acknowledgments

We thank the fermentation unit of the structural biology STP at the Francis Crick Institute. We thank Allison McClure for stimulating discussions. This work has received funding from the European Union’s Horizon 2020 research and innovation programme under the Marie Skłodowska-Curie (895786 to BC), Wellcome Trust Senior Investigator Awards (106252/Z/14/Z and 219527/Z/19/Z to JFXD), European Research Council Advanced Grant (669424-CHROMOREP to JFXD) and the Boehringer Ingelheim Fonds (to GCL). This work was supported by the Francis Crick Institute, which receives its core funding from Cancer Research UK (FC001066), the UK Medical Research Council (FC001066), and the Wellcome Trust (FC001066). For the purpose of Open Access, the author has applied a CC BY public copyright licence to any Author Accepted Manuscript version arising from this submission.

## Author contributions

Conceptualization: BC, APB, JFXD

Methodology: BC, APB, JFXD

Investigation: BC

Resources: BC, GCL, MM

Writing – original draft: BC, APB, JFXD

Writing – review & editing: BC, APB, JFXD

Supervision: JFXD

Project administration: BC, APB, JFXD

Funding acquisition: BC, JFXD

## Supplementary figure legends

**Figure S1. Related to Figure 1.**
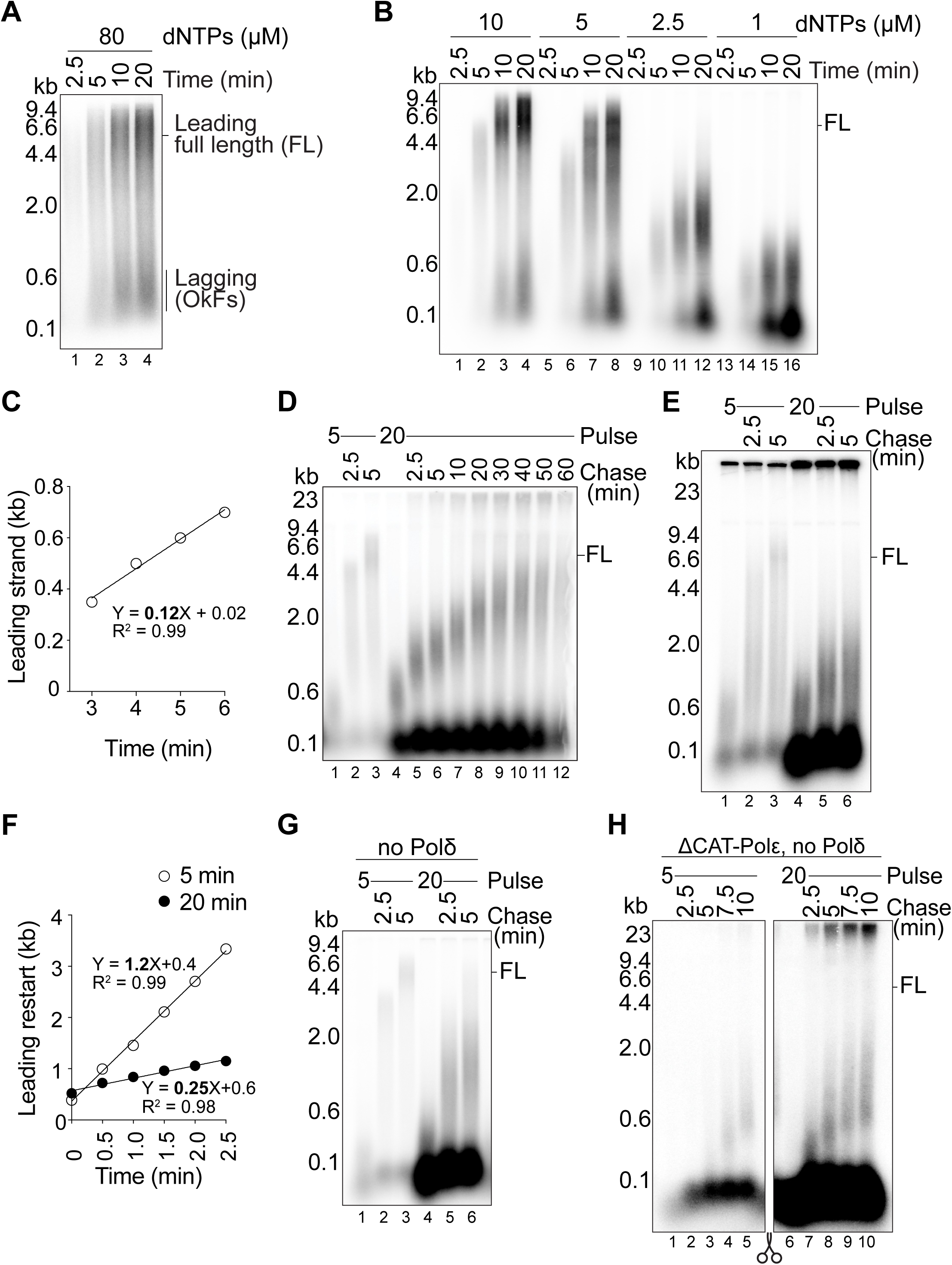
**(A)** Time course of DNA replication reaction with 80 μM each dNTP. **(B)** DNA replication reaction for 2.5, 5, 10 and 20 min in the presence of 10, 5, 2.5 or 1 μM each dNTP. We chose 1 μM dNTP as our ‘low dNTP’ concentration as it still allows to visualise leading and lagging strands, while leading strand is short enough to evaluate restart. **(C)** Quantification of bulk leading strand DNA synthesis speed from 3 to 6 min in Figure 1A. Calculated speed of leading strand synthesis at low dNTP concentration is 0.12 kb/min. **(D)** Pulse-chase DNA replication reactions. 5 and 20 min low dNTP pulses chased for 2.5 and 5 min and 20 min pulse further chased every 10 min up to 60 min. **(E)** Pulse-chase DNA replication reactions denatured with formaldehyde/formamide (See DNA replication reaction with purified proteins in Materials and Methods for details). **(F)** Quantification of bulk leading strand DNA synthesis restart after 5 and 20 min low dNTP pulses from Figure 1D. Calculated speed of leading strand restart is 1.2 kb/min after 5 min and 0.25 kb/min after 20 min. **(G)** Pulse-chase DNA replication reactions omitting Polδ from all reactions. **(H)** Pulse-chase DNA replication reactions where Polε was substituted with the non-catalytic mutant (ΔCAT) Polε, and Polδ was omitted. Polα is the only active polymerase in these reactions. Both gel images in this figure were cropped from a common gel and are comparable since they represent the same exposure of the common gel image. (FL) represents Full leading strand Length.

**Figure S2. Related to Figure 2.**
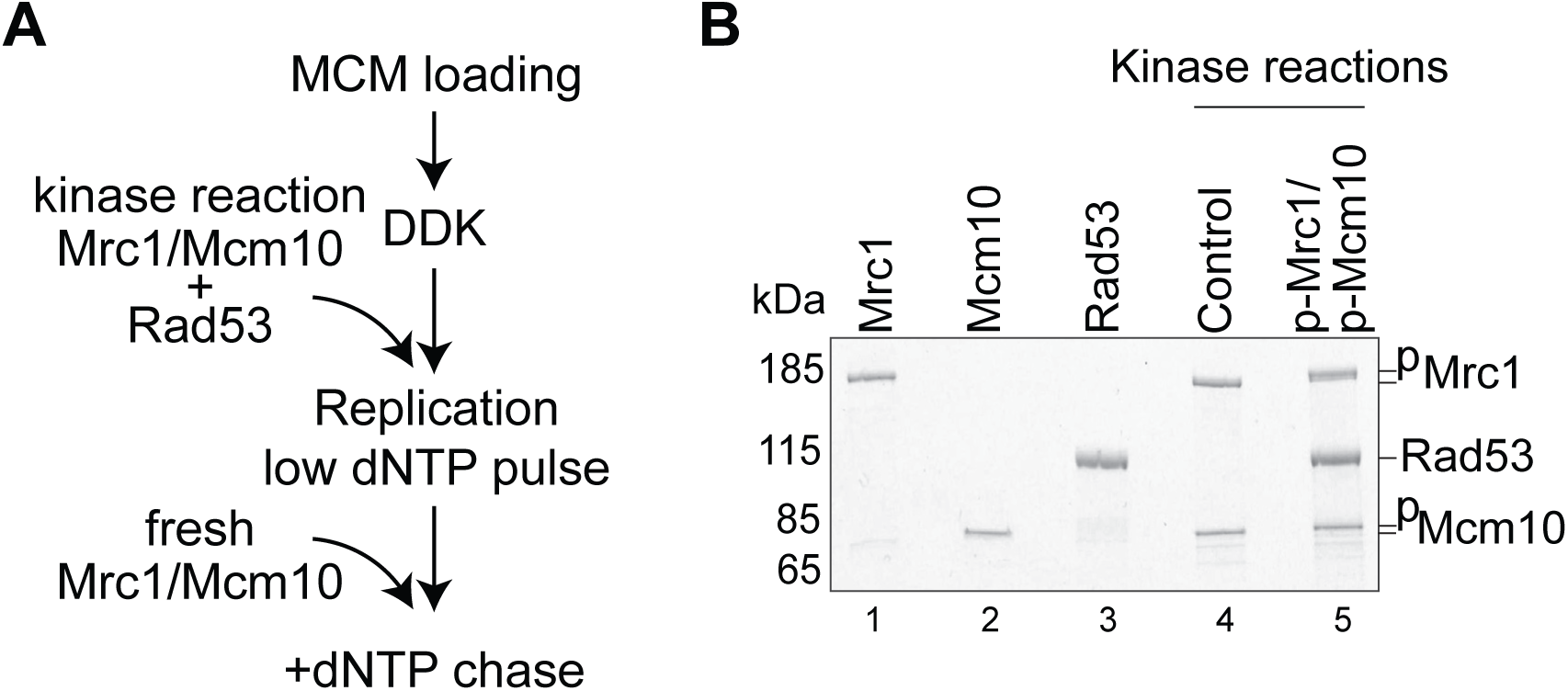
**(A)** Schematic of the experimental steps in Figure 2B. Kinase reactions with Mrc1, Mcm10 and buffer or kinase Rad53 for 30 min were added to DNA replication reactions at low dNTP concentration (pulse) for 5 or 20 min. 30 seconds before adding dNTP chase, fresh Mrc1/Mcm10 were added to all pulses. **(B)** SDS-PAGE and Coomassie staining of individual purified proteins next to 5 μl of the kinase reactions used in Figure 2B (Control and p-Mrc1/p-Mcm10).

**Figure S3. Related to Figure 3.**
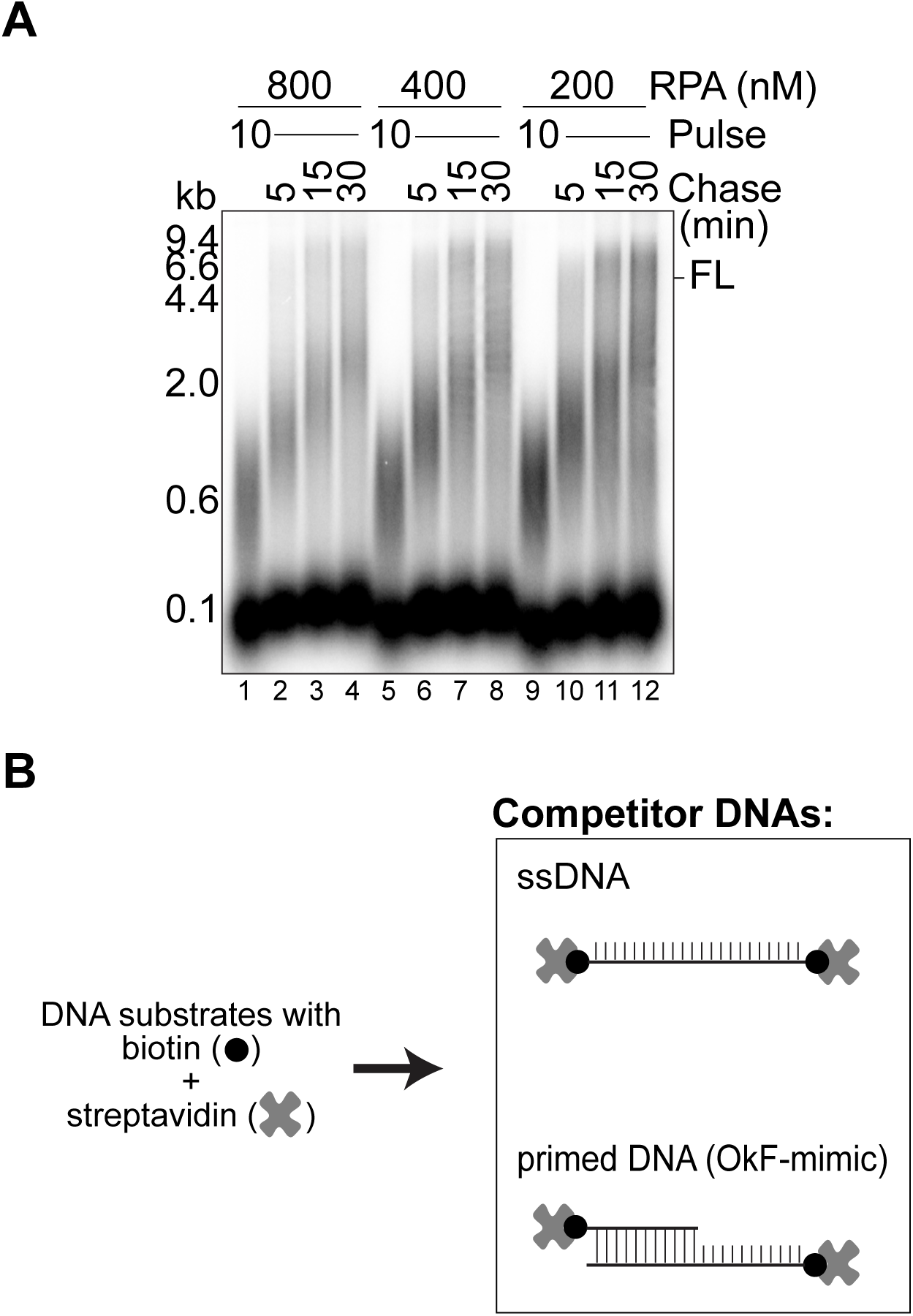
**(A)** Pulse-chase DNA replication reactions. 10 min low dNTP pulses with 800, 400 or 200 nM RPA, chased each for 5, 15 and 30 min. **(B)** Scheme of the competitor ssDNA and competitor primed DNA. DNA oligonucleotides labelled with biotin were incubated with streptavidin to block the ends. (FL) represents Full leading strand Length.

**Figure S4. Related to Figure 4.**
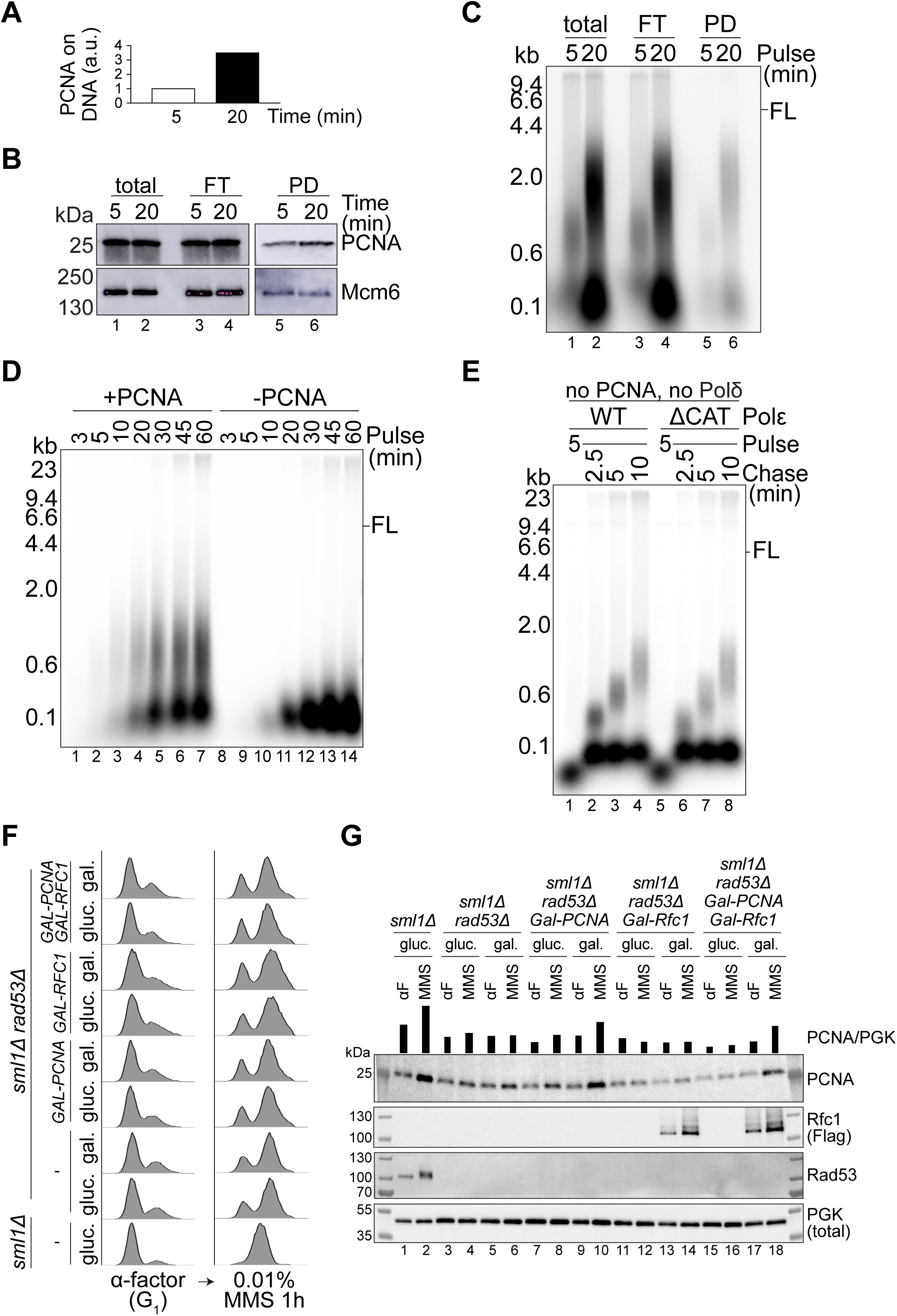
**(A)** Quantification of levels of PCNA loaded on DNA at 20 min. Graph represents immunoprecipitated PCNA levels at 20 min relative to 5 min, each normalised to the Mcm6 loading control. **(B)** Western blot of PCNA and Mcm6 proteins in total, flow through (FT), and pull down (PD) samples from the immunoprecipitation of replicated (biotinylated) DNA in **Figure S4A**. **(C)** Autoradiograph of replicated DNA in total, FT and PD samples from **Figure S4A**. **(D)** DNA replication at low dNTP concentration in the presence or absence of 25 nM PCNA. **(E)** Pulse-chase DNA replication reactions with WT or ΔCAT Polε. PCNA and Polδ were omitted. In lanes 5 to 8, Polα is the only active DNA polymerase present in the reaction. **(F)** Flow cytometry analysis of samples from experiment in Figure 4F. Each exponentially growing strain was synchronised in α-factor (left column) and then released for 1h into medium containing 0.01% MMS (right column) in the specified yeast extract peptone (YP) media supplemented with 2% glucose (gluc.) or galactose (gal.). **(G)** Western blot of whole cell protein extracts from experiment in Figure 4F. Shown are PCNA, Rfc1 (Flag), Rad53 and PGK (total). (FL) represents Full leading strand Length.

**Figure S5. Related to Figure 5.**
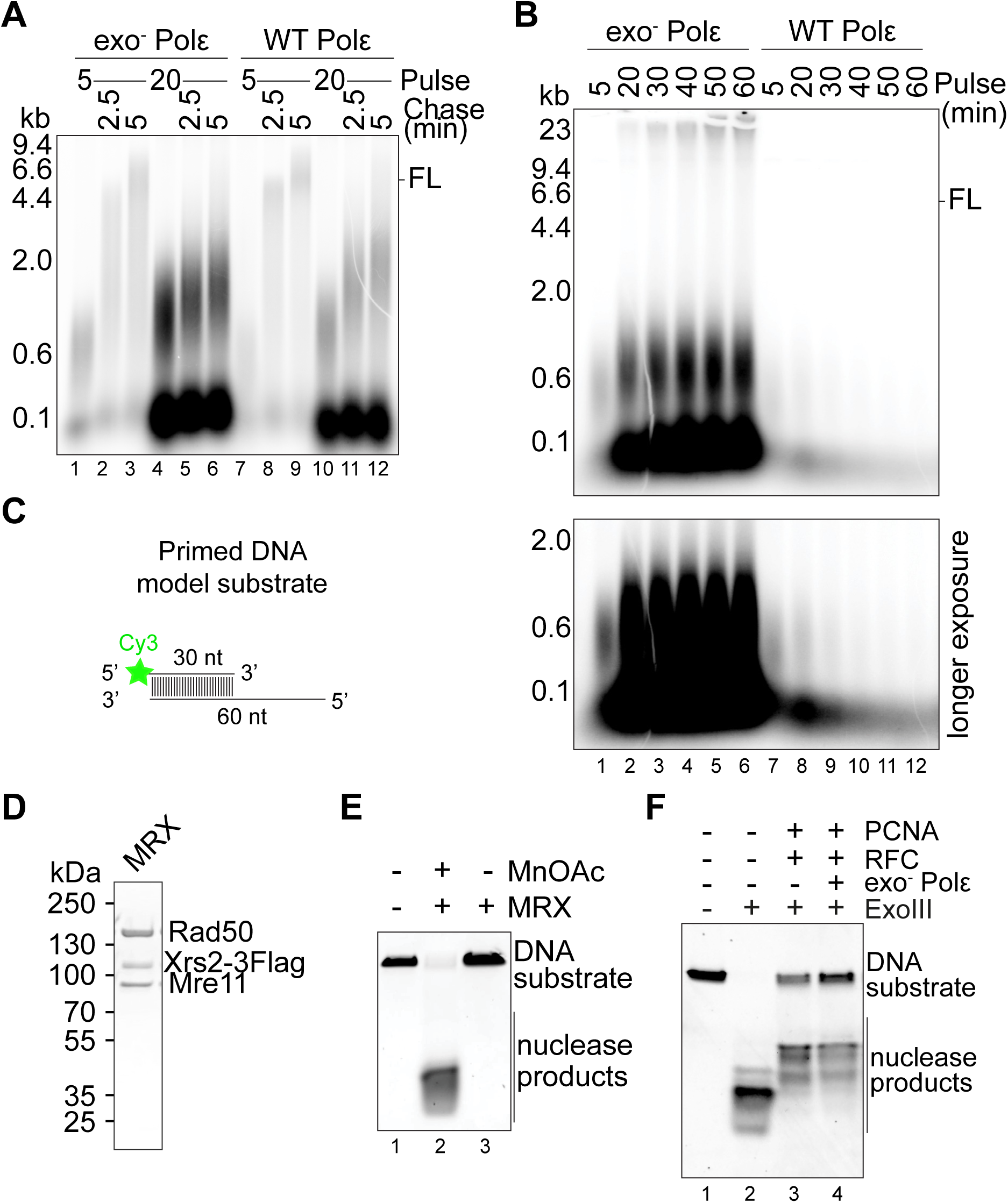
**(A)** Pulse-chase DNA replication reactions. 5 and 20 min low dNTP pulses chased each for 2.5 and 5 min, containing either exo^−^ or WT Polε. **(B)** Time course of DNA replication reactions up to 60 min always in low dNTP concentrations containing either exo^−^ or WT Polε. **(C)** Scheme of the primed DNA model substrate used in nuclease reactions. 30 nt oligonucleotide labelled with fluorescent Cy3 at 5’ is annealed to a 60 nt oligonucleotide creating a 3’ ds/ssDNA junction. **(D)** Purified recombinant budding yeast Mre11, Rad50, Xrs2 (MRX) complex analysed by SDS–PAGE with Coomassie staining. **(E)** Nuclease reactions with MRX nuclease in the presence or absence of 2 mM manganese acetate (MnOAc) to confirm specificity of nuclease activity of our purified MRX complex separated in TBE-PAGE and imaged for Cy3 fluorescence. **(F)** Nuclease reactions performed for 15 min with 2U ExoIII nuclease in the absence or presence of 250 nM PCNA, 125 nM RFC and 100 nM exo^−^ Polε separated in TBE-PAGE and imaged for Cy3 fluorescence. Longer reaction times unravel a role for exo^−^ Polε also in protection of DNA ends from nucleolytic attack. (FL) represents Full leading strand Length.

**Figure S6. Related to Figure 4.**
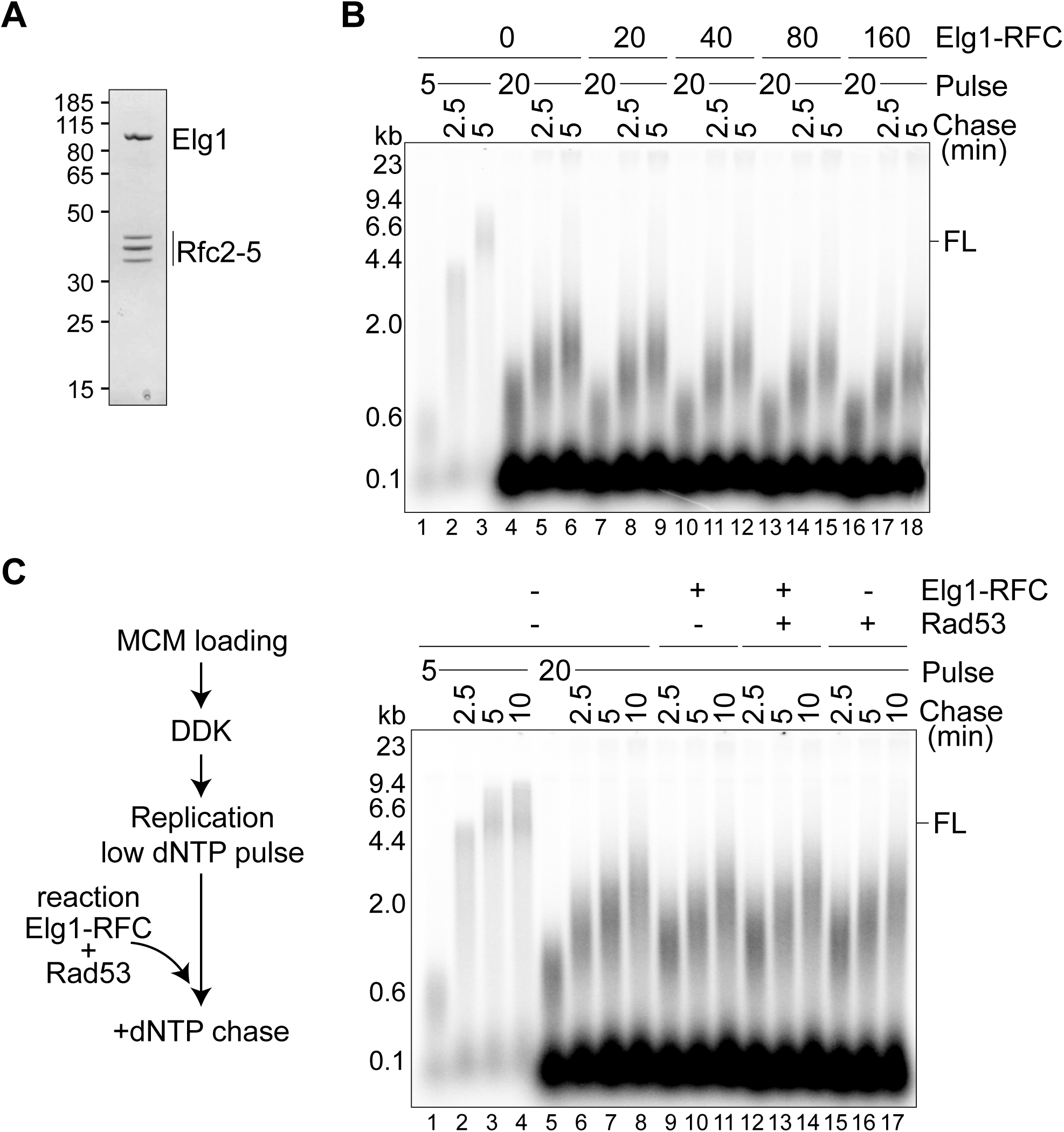
**(A)** Purified recombinant budding yeast Elg1-Rfc2-5 complex by SDS–PAGE with Coomassie staining. **(B)** Pulse-chase DNA replication reactions. 5 or 20 min low dNTP pulses with 0, 20, 40, 80, 160 nM Elg1-Rfc2-5, chased each for 2.5 and 5 min. **(C)** Pulse-chase DNA replication reactions. 5 or 20 min low dNTP pulses. Reactions containing combinations of Elg1-Rfc2-5 and Rad53 or their buffers (see Rad53 kinase reaction in Materials and Methods for details) were added to each pulse 30 sec before chasing with dNTPs for 2.5, 5 and 10 min. (FL) represents Full leading strand Length.

